# Single-cell assessment of trophoblast stem cell-based organoids as human placenta-modeling platforms

**DOI:** 10.1101/2022.11.02.514970

**Authors:** Matthew J. Shannon, Gina L. McNeill, Burak Koksal, Jennet Baltayeva, Jasmin Wächter, Barbara Castellana, Maria S. Peñaherrera, Wendy P. Robinson, Peter C. K. Leung, Alexander G. Beristain

**Author notes:** To whom correspondence should be addressed: Alexander G. Beristain, The British Columbia Children’s Hospital Research Institute, The University of British Columbia, Vancouver, British Columbia, Canada. V5Z 4H4.

## Abstract

The recent discovery of human trophoblast stem cells (hTSC) and techniques allowing for trophoblast organoid (TOrg) culture have established promising approaches for studying human trophoblast development. To validate the accuracy of these models at single-cell resolution, we directly compared *in vitro* TOrg cultures derived from primary progenitor cytotrophoblasts (CTB) or commercially available hTSC lines to *in vivo* human trophoblasts using a scRNA-seq approach. While patient-derived (PD)- and hTSC-derived TOrgs overall reflect cell differentiation trajectories with accuracy, specific features related to trophoblast state make-up, distinct sub-paths of differentiation, and predicted transcriptional drivers regulating stem cell maintenance were shown to be misaligned in the *in vitro* platforms. This is best exemplified by the identification of a distinct progenitor state in hTSC-derived TOrgs that showed characteristics of CTB- and extravillous-like cell states. Together, this work provides a comprehensive resource that identifies underlying strengths and limitations of current TOrg platforms.

**Summary Statement:** Single-cell transcriptomics provides comprehensive comparison between trophoblast organoids derived from commercially available trophoblast stem cells and first-trimester primary human cytotrophoblasts.

**HIGHLIGHTS:**

- An integrated single cell transcriptomic atlas of placental and organoid trophoblasts establishes a comprehensive and public web-based resource
- Direct comparison of trophoblasts from placental/decidual tissue to trophoblasts extracted from two distinct organoid platforms highlights both conserved and divergent features
- Computational modeling describes novel trophoblast states and routes of cell differentiation in human trophoblast organoids

**IN BRIEF:** While the merits and utility of current trophoblast organoid cultures have been established, high-resolution assessment and comparison of conserved and divergent features of these systems to cell states and differentiation trajectories of trophoblasts *in situ* or *in vitro* has not been performed. Here, Shannon et al. generate a single-cell transcriptomic atlas of two trophoblast organoids that comprehensively define the similarities and discrepancies in relation to trophoblasts from the placental-maternal interface.

## INTRODUCTION

In mammalian development, the placenta forms the mechanical and physiological link between maternal and fetal blood. Specialized placental cells called trophoblasts, derived from the trophectoderm of the implanting blastocyst, perform many of the placenta’s transport, endocrine, and immunogenic functions^1, 2^. In rodents and primates that have haemochorial placentas (i.e., trophoblasts are subject to direct contact with maternal blood), nutrient and oxygen transfer between maternal and fetal circulations are controlled in part by the actions of syncytium-forming and invasive trophoblast subsets^3, 4^. While trophoblasts perform similar functions across species, trophoblast developmental trajectories in rodents and humans are ultimately different. These differences are underlined by distinct species-specific molecular programs controlling trophoblast stem cell maintenance and differentiation^5–8^. Because of these molecular and cellular distinctions, significant limitations exist when using rodent models to study human placental development. At the same time, major ethical constraints, difficulties in accessing human placental tissue for research, and sub-optimal *in vitro* systems have limited our progress in understanding the cellular processes that are important in human placentation.

In 2018, three separate reports described the establishment of long-term, two-dimensional (2D)^8^ and three-dimensional (3D)^9, 10^ regenerative stem cell-based trophoblast cultures. Derived from progenitor cytotrophoblasts (CTB) of early first-trimester chorionic villi, 2D human trophoblast stem cell (hTSC) and 3D trophoblast organoid (TOrg) cultures can be perpetuated long-term in the presence of Wingless/Integrated (WNT) and EGF signals concurrent with TGFβ receptor and Rho-associated protein kinase (ROCK) inhibition^8–10^. Both hTSC-TOrgs and freshly-isolated patient-derived (PD)-TOrgs spontaneously fuse to generate hormone-producing multinucleated syncytiotrophoblast (SCT). Removal of WNT potentiation, in combination with NRG1 stimulation, promotes CTB to differentiate along the extravillous pathway, generating HLA-G-expressing extravillous trophoblasts (EVT)^9, 10^. These newly described trophoblast models establish state-of-the-art systems to study human trophoblast development.

Recently, a study comparing blastocyst- and CTB-derived hTSC lines^8^ to primary CTB-derived TOrgs identified a number of molecular features unique to hTSCs that are absent in TOrgs and chorionic villous CTB^11^. Notably, HLA-typing showed that hTSCs cultured in 2D express detectable levels of surface HLA-A, -B, and -G, where a transition to 3D culture dampened class-I HLA expression^11^. Moreover, bulk RNA-seq analyses showed that hTSCs and TOrgs cluster distinct from each other, suggesting that major transcriptomic differences and/or differences in cellular composition exist between these models. Importantly, while founding hTSC^8^ and TOrg^9, 10^ studies have primarily focused on their similarities to trophoblasts *in vivo*, discrepancies between TOrgs and *in vivo* trophoblasts have not been directly explored or defined. Therefore, a more thorough evaluation of current stem cell-based trophoblast cultures is warranted.

Here, previously reported hTSC-TOrg^12^ and in-house generated PD-TOrg single-cell RNA-sequencing datasets were used to comprehensively compare TOrg cell states and transcriptomics. These datasets were also compared to corresponding/equivalent trophoblasts from first-trimester chorionic villi and decidua using publicly available scRNA-seq datasets^12, 13^. These new data are made available at: https://humantrophoblastmodels.shinyapps.io/mjs_trophoblasts_2022/, providing a rich resource for direct transcriptomic comparison between TOrgs and trophoblasts residing within the maternal-placental interface. While major trophoblast states are generally conserved in hTSC- and PD-TOrgs, cluster analyses identified an expanded EVT progenitor-like state in hTSC-TOrgs that is modestly present in PD-TOrgs and minimally present *in vivo*. Analyses examining cell-ligand interactions, transcription factors (TFs) driving trophoblast state-specific regulons, and trophoblast developmental trajectories further highlights important differences between these *in vitro* and *in vivo* systems.

## RESULTS

### Generation of a PD-TOrg single-cell transcriptomic resource and inter-model comparison

Because a PD-TOrg single-cell transcriptomic resource has not yet been established, we first set out to generate PD-TOrgs and characterize general features of this primary 3D model in tandem with hTSC-TOrgs prior to high-dimensional transcriptomic workflow. When embedded in Matrigel and cultured in the presence of WNT- and EGF-potentiating factors (herein referred to as Wnt^+^ media), both primary CTB and hTSC lines (CT27, CT29, CT30) established 3D structures defined by peripheral layers of CTB and an inner SCT core (Fig. 1A; Fig. S1A-B). All three hTSC lines consistently formed large (>25,000 μm^2^) organoids by day 9 of culture (Fig. 1B). In contrast, assessment of two PD-TOrgs (P1069, P1111) showed that primary CTBs generated equivalently sized structures by day 15 of culture, highlighting a growth advantage that hTSC-lines have over PD-TOrgs (Fig. 1B; Table S1). In line with previous reports^9, 10^, the removal of the WNT-potentiating factors R-spondin and CHIR99021 (herein referred to as Wnt^-^ media) resulted in characteristic EVT-like outgrowths in PD- and hTSC-TOrgs (Fig. S1A). Assessment of gene markers representing CTB (*ITGA6, TEAD4*), SCT (*ERVFRD-1, CGB*), and immature (*ITGA5, NOTCH1*) and mature (*HLA-G, ITGA1*) EVT, showed that both types of TOrgs maintained steady-state levels of CTB and SCT genes (Fig. S1C). Both organoid systems further showed expression of mature EVT markers by culture endpoint (Fig. S1C) in addition to HLA-G surface expression (Fig. S1D). DNA methylation levels at 6 CpGs within the *ELF5* promoter in TOrg CTBs as well as in CTB/SCT isolated from first-trimester chorionic villi showed clear hypomethylation of the ELF5 promoter in all trophoblasts (Fig. S1E). These data show that our primary CTB and hTSC-lines establish 3D TOrgs that retain the ability to differentiate into SCT- and EVT-like cells.

**Figure 1:**
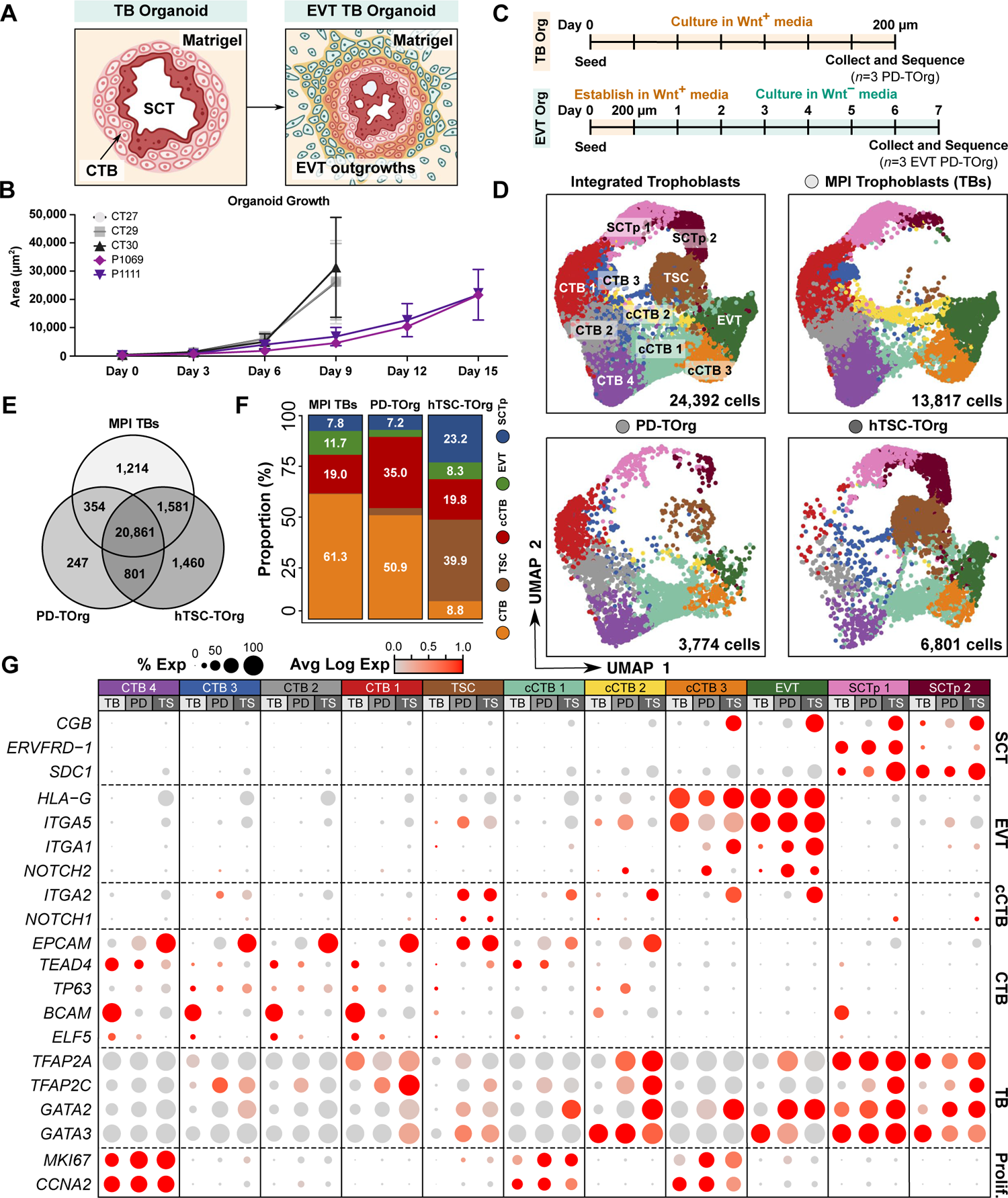
Single-cell characterization of MPI, PD-TOrg, and hTSC-TOrg trophoblasts. **(A)** Schematic demonstrating trophoblast organoid establishment and EVT-differentiation in Matrigel. Cytotrophoblast (CTB), syncytiotrophoblast (SCT), extravillous trophoblast (EVT). **(B)** Growth curve of PD-TOrgs (P1069, P1111) and hTSC-TOrgs (CT27, CT29, CT30) over 9 and 15 days of culture. Organoid size is measured as the image area (μm^2^) acquired using a 10X objective. Median values are shown and error bars represent standard error. **(C)** Workflow used for generating pooled regenerative and EVT-differentiated PD-TOrg single-cell suspensions. Pooled (*n*=3) regenerative PD-TOrgs were cultured for 7 days in WNT-potentiating media (Wnt^+^) before sequencing. Pooled (*n*=3) EVT-differentiated PD-TOrgs were established to 200 μm diameter in Wnt^+^ media then cultured for 7 days in media lacking R-spondin and CHIR99021 (Wnt^-^) before sequencing. **(D)** UMAP plot of 24,392 integrated; 13,817 MPI; 3,774 PD-, and; 6,801 hTSC-TOrg trophoblasts. 11 clusters representing four trophoblast subtypes were found. MPI: maternal-placental interface, TB: trophoblast, column CTB: cCTB, trophoblast stem cell: TSC, SCT precursor: SCTp. **(E)** Venn diagram comparing bulk MPI, PD-, and hTSC-TOrg transcriptomes. **(F)** Proportion of captured trophoblasts in each binned scRNA-seq-defined state (CTB: CTB1-4; cCTB: cCTB1-3; SCTp: SCTp1-2) in each model. **(G)** Dot plot showing the percent (0-100%) and average expression level (value of 0.0 to 1.0) of known trophoblast and proliferation gene markers across each cluster and in each model. PD: PD-TOrg; TS: hTSC-TOrg.

Having established both regenerative and EVT-differentiated PD-TOrg cultures, PD-TOrg single-cell cDNA libraries were generated from three distinct first trimester chorionic villi CTB preparations (*n*=3) (Table S2): Regenerative TOrgs were cultured to 200 μm in Wnt^+^ media while EVT-differentiated TOrgs were first established to 200 μm diameter in Wnt^+^ media followed by 7-days culture in EVT-differentiating Wnt^-^ media; this approach is identical to that previously applied in generating single-cell cDNA libraries in hTSC-TOrgs^12^. TOrg cells were then dissociated and pooled to generate cDNA libraries following 10x Genomics Chromium workflow (Fig. 1C). This dataset was merged with published *in vivo* human trophoblast^12, 13^ and hTSC-TOrg^12^ single-cell transcriptomic atlases. Clinical characteristics and sequencing metrics for each patient and sample are summarized (Table S2A). Following preprocessing and filtering of the integrated data, 24,392 trophoblasts from chorionic villi/decidua and TOrgs were identified by expression of trophoblast lineage genes and the absence of non-trophoblast lineage markers.

Eleven trophoblast clusters were resolved representing four CTB-like states (CTB1-4; *TEAD4*^+^, *ELF5^+^*, and *EGFR^+^*), two SCT precursors (SCTp) expressing SCT-associated genes (SCTp1-2; *ERVFRD-1^+^*, *CGB^+^*), three column CTB-like states (cCTB1-3; *NOTCH1^+^*, *ITGA5^+^*), one EVT-like state (EVT; *HLA-G^+^*, *ITGA1^+^*), and one state predominant in hTSC-TOrgs that we termed TSC (Fig. 1D; Table S3A). Transcriptomic similarity between maternal-placental interface (MPI; *in vivo*) and TOrg trophoblasts, assessed using Pearson’s correlation coefficient (PCC) analysis, showed PD-TOrg and hTSC-TOrg signatures to be 92.9% and 79.5% similar to *in vivo* trophoblasts, respectively. Exploring this further, 20,861 genes were common to *in vivo* trophoblasts and TOrgs while 1214, 247, and 1460 genes were unique to MPI, PD-, and hTSC-TOrg trophoblasts, respectively (Fig. 1E; Table S4A).

*In vivo,* 61% of trophoblasts were progenitor CTB whereas column-like, EVT-like, and SCT precursors made up 19%, 12%, and 8% of all trophoblasts, respectively (Fig. 1F). Comparing TOrg and *in vivo* trophoblast-subtype proportions showed that PD-TOrgs better reflect *in vivo* trophoblast composition than hTSC-TOrgs (Fig. 1F; Table S5). For example, PD-TOrgs, similar to *in vivo* trophoblasts, were dominated by progenitor-like CTBs (51% of cells) (Fig. 1F). This contrasted hTSC-TOrgs, where CTB contributed to less than 9% of cells (Fig. 1F). Notably, in hTSC-TOrgs the TSC state predominated, comprising 40% of all cells (Fig. 1D; 1F). By comparison, the TSC state represented 3% of cells in PD-TOrgs and 0.3% *in vivo* (Fig. 1F). Other notable differences include increased frequencies of column-like cells in PD-TOrgs and SCTp in hTSC-TOrgs (Fig. S1C).

Comparing the expression of trophoblast lineage and proliferation-associated genes showed that both types of TOrgs resemble *in vivo* transcriptomic patterns (Fig. 1G). For example, CTB4, cCTB1, and cCTB3 states across TOrg and MPI trophoblasts showed higher levels of the proliferative genes *MKI67* and *CCNA2* (Fig. 1G). Moreover, states representing cell differentiation endpoints (i.e., SCTp1, SCTp2, EVT) showed alignment of key transcripts, including the fusogenic gene *ERVFRD-1* in SCTp1 and the extravillous markers *HLA-G* and *ITGA5* in EVT (Fig. 1G). To benchmark the unique TSC state, we compared trophoblast subtype transcriptomic signatures independent of model using a PCC analysis. The TSC state was 87% similar to EVT, 81% similar to cCTB, 73% similar to CTB, and 69% similar to SCTp, suggesting that the TSC state harbours both progenitor-like and differentiated features (Fig. S1F).

In summary, scRNA-seq showed that PD- and hTSC-TOrgs have conserved, as well as distinct, transcriptomic features when compared to *in vivo* trophoblasts. Overall, PD-TOrgs better recapitulate the MPI trophoblast landscape over hTSC-TOrgs from the perspectives of global gene expression, cell-state heterogeneity, and specific trophoblast gene marker expression.

### Defining ligand-receptor cell communication networks in organoid and *in vivo* trophoblasts

Cell-cell signaling networks influence tissue functionality^14, 15^. Therefore, when modeling trophoblasts *in vitro* it is important to assess how accurately models reproduce specific cellular and molecular niches. To do this, MPI, PD-, and hTSC-TOrg trophoblasts were individually re-clustered and model-specific trophoblast states were binned into CTB, TSC, cCTB, EVT, and SCTp groups (Fig. 2A). Using the Connectome package^16^, cell-cell, cell-matrix, and cell-signaling topologies were visualized (Fig. 2B). We assessed the Hippo^17^, BMP^18^, WNT^19^, VEGF^20, 21^, NOTCH^19^, metalloprotease/extracellular matrix^22, 23^, and EGF^24^ gene families because of their importance in regulating trophoblast renewal, differentiation, and function. Centrality analysis assessed which trophoblast-subtype predominantly produced ligands and which subtype expressed receptors for those ligands (correlated receivers). *In vivo*, EVT showed the greatest ligand production potential with the exceptions of laminin (predominantly SCTp) and NOTCH (highest in cCTB) (Fig. 2B). By comparison, PD-TOrg ligand production was predominated by TSC and EVT states, with the exceptions of EGF and WNT (highest in SCTp) and laminin (predominantly CTB). hTSC-TOrg ligand production varied, with the CTB state driving matrix glycoprotein ligand production; the cCTB state driving ADAM, EGF, assorted matrix, and WNT ligand production; the EVT state driving laminin and VEGF ligand production; the SCTp state driving BMP and NOTCH ligand production, and no significant MMP ligand production determined (Fig. 2B). Regarding correlated receivers, *in vivo* cCTB and EVT states and PD-TOrg TSC, cCTB, and EVT states showed strong receptor conservation, highlighting broad enrichment of metalloprotease (ADAM, MMP), matrix, EGF, NOTCH, and VEGF categories (Fig. 2B). In contrast, hTSC-TOrgs demonstrated markedly different patterns in receptor/receiver profiles compared to *in vivo* trophoblasts and PD-TOrgs, showing reduction in EGF, laminin, matrix, MMP, and WNT receptor signals (Fig. 2B).

**Figure 2:**
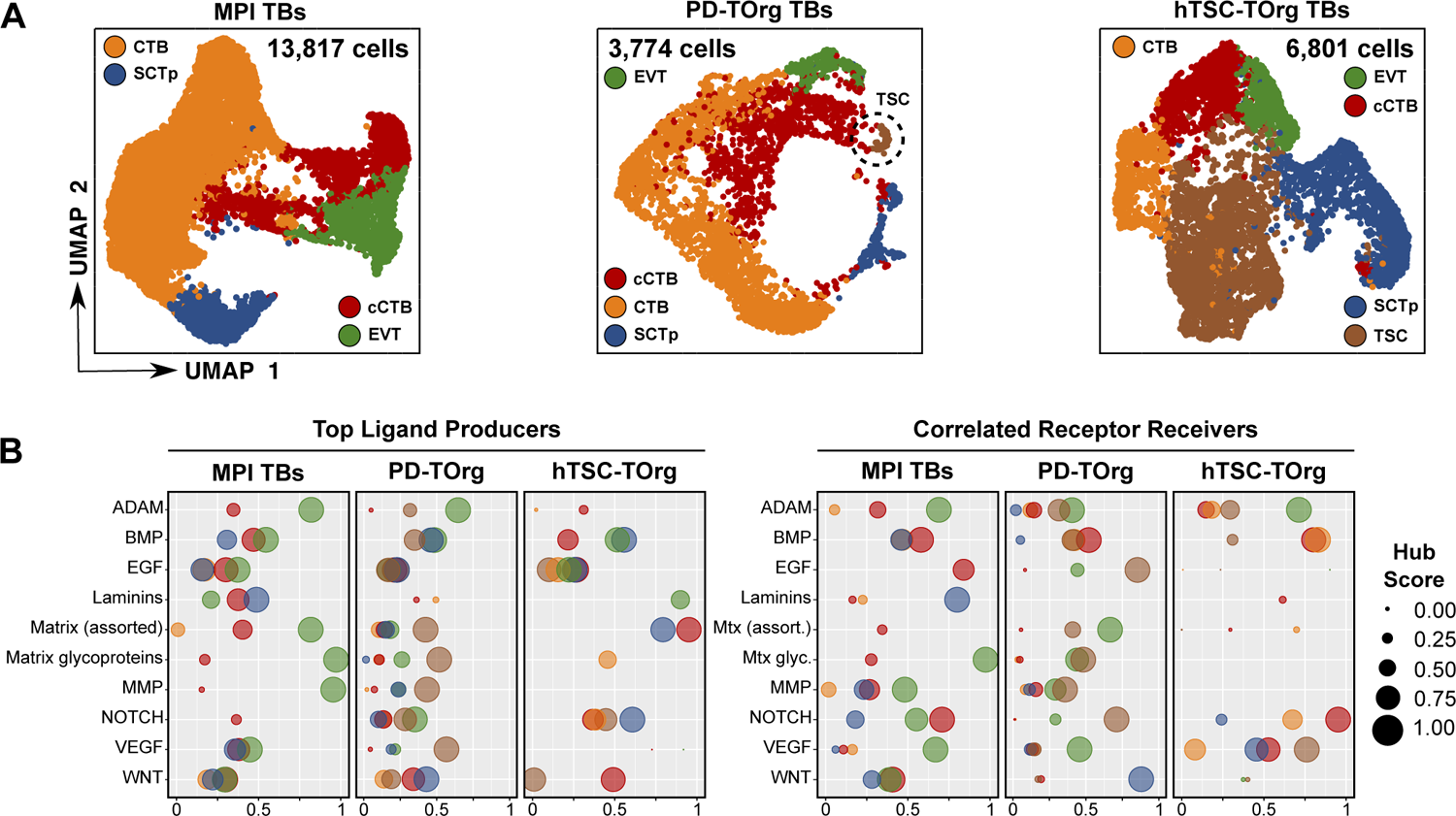
Comparing *in vivo* TB and TOrg cell-cell signalling networks. **(A)** Re-clustered MPI, PD-, and hTSC-TOrg UMAP plots. Trophoblast states are denoted by the color legend **(B)** Dot plots summarizing the ligand producers and subtypes expressing correlated receptors of ADAM, BMP, EGF, Laminin, Matrix/Matrix glycoprotein, MMP, NOTCH, VEGF, and WNT signaling families in MPI, PD-, and hTSC-TOrg trophoblasts.

### hTSC-TOrgs deviate from initial *in vivo* and PD-TOrg progenitor trajectories but reproduce mature trophoblast endpoints

To examine how TOrgs recapitulate *in vivo* CTB differentiation, Monocle2 pseudotime ordering was performed^25, 26^ (Fig. 3A). As described previously^12, 13^, pseudotime ordering of *in vivo* trophoblasts predicts a CTB state origin that develops toward EVT or SCTp states (Fig. 3A). As *in vivo* CTB differentiate along the extravillous branch, expected gene expression changes are observed. This is exemplified by decreases in progenitor-associated transcripts, transient increases in column-associated genes (*NOTCH1, ITGA2*), and endpoint expression of mature EVT genes (*NOTCH2, ITGA1*) (Fig. 3B). Expected expression kinetics were also observed along the villous branch as CTB differentiate into the SCTp state and begin to express the fusogenic genes *ERVW-1* and *ERVFRD-1* (Fig. 3B). Villous and extravillous trajectories, and associated gene expression kinetics were mostly conserved in PD-TOrgs, although the transition from CTB to cCTB states was less distinct (Fig. 3A). Further, differences in *CGB*, *ITGA5*, and *ITGA1* kinetics were observed, where levels increased at the tail ends of both SCT (*ITGA5, ITGA1*) and EVT (*CGB*) pathways (Fig. 3B). Compared to pseudotime ordering of MPI cells, hTSC-TOrgs showed greater deviances in cell trajectory and gene expression alignment. Firstly, the trajectory start-point of hTSC-TOrgs consisted of TSC state cells; this is in contrast to the *in vivo* and PD-TOrg origins populated predominately by CTB state cells (Fig. 3A). Secondly, direct differentiation from the TSC state into SCTp was observed along the villous branch, while along the extravillous branch, the TSC state preceded establishment of the CTB state, which subsequently transitioned into the cCTB then EVT states (Fig. 3A). Lastly, transient upregulation of *NOTCH1* and *ITGA2* midway through EVT differentiation was not observed in hTSC-TOrgs, though expected endpoint SCTp and EVT gene signatures were retained (Fig. 3B).

**Figure 3:**
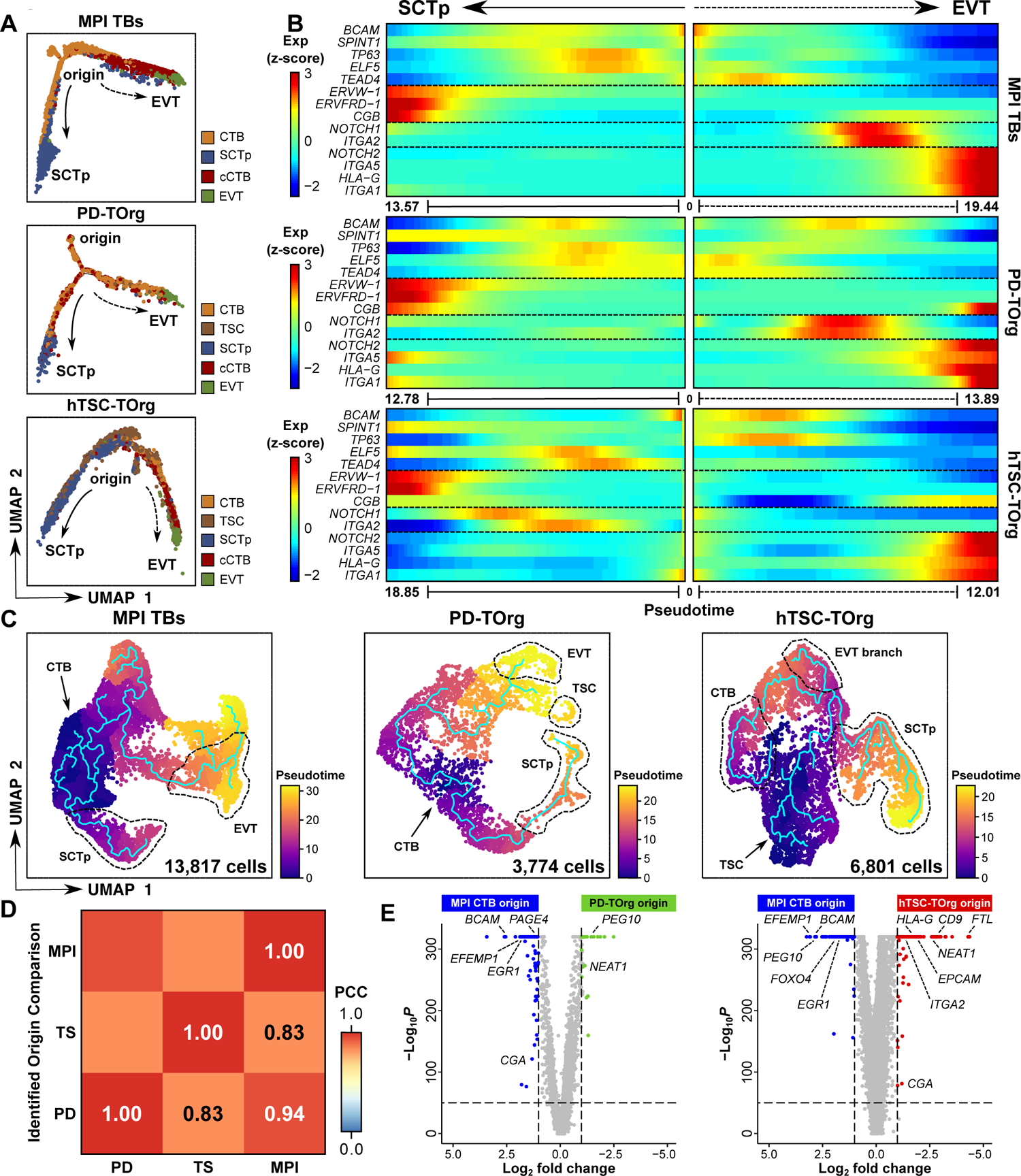
Assessing *in vivo* trophoblast and TOrg differentiation trajectories. **(A)** Monocle2 pseudotime ordering of MPI, PD-, and hTSC-TOrg trophoblasts along the villous and extravillous paths of differentiation. Trophoblast states (CTB, SCTp, cCTB, EVT, TSC) are indicated by the color legend. **(B)** Branched heatmap showing gene expression changes across inferred pseudotime towards SCTp and EVT endpoints; pseudotime origin (0) and endpoint values are shown under heatmaps. **(C)** MPI, PD-, and hTSC-TOrg UMAP plots overlain with constructed Monocle3 lineage trajectories. Cells are color coordinated by relative pseudotime value. Select trophoblast states/differentiation branches are encircled. **(D)** Pearson’s correlation coefficient heatmap between MPI, PD-, and hTSC-TOrg origins. Model-specific origins were treated as pseudo-bulk samples during correlation and abbreviations are as in Fig. 1. **(E)** Volcano plot showing DGEs between MPI trophoblast origin and PD-TOrg origin (left) and hTSC-TOrg origin (right).

These changes in hTSC-TOrg gene patterns across pseudotime could reflect an hTSC-specific *in vitro* differentiation artifact but may also result from limitations intrinsic to Monocle2, a pseudotime tool that infers simple branched trajectories. We therefore compared model-specific progenitor gene expression signatures (Fig. S2A) and applied Monocle3^27^ trajectory inference to refine trajectory predictions. Monocle3 recapitulated expected *in vivo* cell-state ordering with the addition of a looped trajectory in CTB, suggesting that CTB states are both transient and cyclical (Fig. 3C). Similarly, Monocle3 ordering of PD-TOrgs defined a CTB state origin from which trophoblasts progressed toward either SCTp or EVT states (Fig. 3C). In hTSC-TOrgs, Monocle3 resolved three developmental trajectories originating from the unique TSC state: one branch developing toward the EVT endpoint, another toward SCTp, and a third toward the CTB state (Fig. 3C). This altered hTSC-TOrg trajectory highlights a limitation for confining hTSC-TOrg development along two differentiation paths and suggests that substantial differentiation complexity exists *in vitro*.

Because the hTSC-TOrg-predicted origin differed from the *in vivo* and PD-TOrg CTB origin, we next set out to compare origin-specific transcriptomic signatures between *in vivo* trophoblasts and TOrgs. To do this, we subset cells overlapping with the predicted origin in each model (Fig. S2B) and assessed the transcriptomic similarity of each origin. PCC analysis showed that PD-TOrg origin signatures were 94% similar while hTSC-TOrg origin signatures were 83% similar to origin cells *in vivo* (Fig. 3D). Next, we conducted a DGE between each origin to identify significantly enriched genes in each system. Compared to TOrgs, *in vivo* origin trophoblasts showed elevated levels of *BCAM*, *EGR1*, and *FOXO4* (Fig. 3E; Table S6A). This was in contrast to the PD-TOrg origin that showed enrichment of *NEAT1* transcripts and the hTSC-TOrg origin showing enrichment of *HLA-G*, *ITGA2*, *EPCAM*, and *NEAT1* transcripts (Fig. 3E; Table S6B). This hTSC-TOrg signature suggests a progenitor state that is more differentiated than the origin-aligned cells found *in vivo* and in PD-TOrgs.

### *In vivo* driver transcription factors are conserved in TOrg CTB-like states

Because trajectory modeling suggests that hTSC-TOrg CTB do not function as upstream progenitor cells, we next set out to compare transcriptomic signatures and networks between *in vivo* and TOrg CTBs. Following sub-setting and re-clustering of CTB states (Fig. 4A), transcripts were largely conserved between *in vivo* and TOrg CTBs (Fig. 4A). Specifically, 17,235 genes showed overlap whereas 2551; 681; and 330 genes were unique to MPI, PD-, and hTSC-TOrg CTB, respectively (Fig. S3A; Table S4C). Further, hTSC-TOrg CTBs retained increased levels of EVT-(*ITGA5*, *HLA-G*) and differentiation-associated *GCM1*^28^ transcripts (Fig. S3B). Next, TFs driving CTB-specific regulons were determined using the SCENIC^29^ gene regulatory network inference tool. SCENIC identified 237 *in vivo* CTB regulons, while 253 and 268 regulons were identified in PD- and hTSC-TOrg CTBs, respectively (Table S7A-C). Regulons were ranked based on activity level in CTBs in each model with *SRF* driving the most active *in vivo* regulon and *SMARCA4* and *ETV5* driving the most active PD- and hTSC-TOrg regulons, respectively (Fig. 4B). TFs driving CTB regulons included genes known to be important in human trophoblast lineage (i.e., *GATA2*/*3*, *TFAP2A*/*C*, *TEAD4*), but also TFs not yet explored in human CTB biology (Table S7). Overall, CTB regulons were conserved across models with variation existing at the level of regulon rank (Table S7A-C). To ensure specificity, we expanded SCENIC analysis to include all trophoblasts, assessed regulon activity across trophoblast states, and through this, identified 243 regulons driving CTB, TSC, cCTB, EVT, and SCTp transcriptomes (Fig. S4A; Table S7D). The most important regulons driving trophoblast state/subtype were identified by binarizing regulon activity and selecting for regulons active in at least 50% of cells (Fig. S4B). This identified 6 CTB-specific regulons controlled by *EGR1*, *E2F4*, *FOXP1*, *FOXO4*, *IRF6*, and *TFAP2B,* as well as 2 CTB-enriched regulons (*PBX1*, *TEAD4*). Across all 6 CTB-specific regulons, TF levels were highest *in vivo* with hTSC-TOrg CTB showing lowest expression (Fig. 4C).

**Figure 4:**
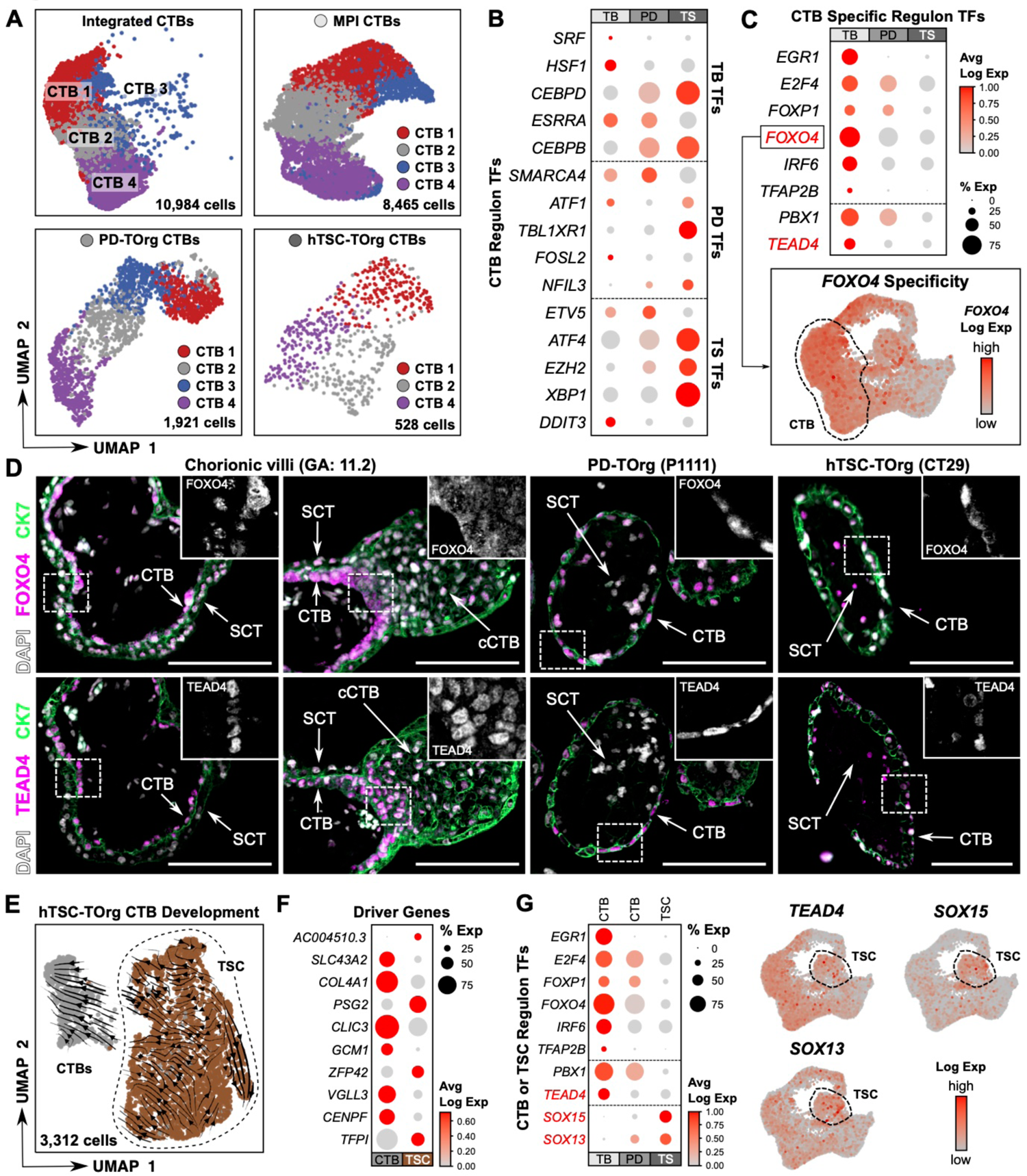
Comparing *in vivo* and TOrg progenitor trophoblast gene expression and regulation. **(A)** Integrated and re-clustered model-specific CTB UMAP plots. **(B)** Dot plot comparing the percent and average expression level of TFs driving the top 5 regulons in MPI (top), PD- (middle), and hTSC-TOrg (bottom) CTB. Legend is shared with panel C. **(C)** Dot plot comparing the percent and average expression level of TFs driving CTB-specific (top) and CTB-enriched (bottom) regulons in *in vivo* and TOrg CTB. *FOXO4* is highlighted with expression (red) plotted in integrated UMAP space. **(D)** Representative immunofluorescence (IF) images of serially sectioned first-trimester chorionic villi (11.2 weeks’ gestation), PD-TOrg (P1111), and hTSC-TOrg (CT29) labelled with antibodies targeting FOXO4 and CK7 (top) and TEAD4 and CK7 (bottom); nuclei are stained with DAPI. Bar = 200 µm. Hashed lines depict enlarged inset of FOXO4 or TEAD4 staining. Cytotrophoblast (CTB), syncytiotrophoblast (SCT), column cytotrophoblast (cCTB) are identified by arrows. **(E)** Re-clustered hTSC-TOrg CTB and TSC UMAP plot. UMAP is overlain with projected RNA velocity vector field. **(F)** Dot plot showing the percent and average expression level of genes driving the TSC and CTB vector field. **(G)** Dot plot comparing the percent and average expression level of TFs driving CTB regulons between *in vivo* CTB, PD-TOrg CTB, and hTSC-TOrg TSC. *TEAD4*, *SOX15*, and *SOX13* are highlighted with expression (red) plotted in integrated UMAP space. TF: transcription factor and abbreviations are as in Fig. 1.

*FOXO4* and *TEAD4* were identified as driver TFs of CTB state regulons *in vivo* and in TOrgs (Fig. 4C). FOXO4 localizes to the SCT of term placentas^30^ but spatial assessment of FOXO4 in the first trimester and the role of FOXO4 in human trophoblast development has not yet been examined. By contrast, the importance of TEAD4 in regulating progenitor CTB renewal and proliferation is more thoroughly understood^31–33^. To verify that these driver TFs localize to CTB/CTB-like cells *in vivo* and within our organoid systems, we examined FOXO4 and TEAD4 localization in chorionic villi and in TOrgs by immunofluorescence microscopy (IF) (Fig. 4D). In chorionic villi, strong nuclear FOXO4 and TEAD4 signals were present in CTB (Fig. 4D). FOXO4 signal was additionally present in the nuclei of syncytiotrophoblast, though this signal was less intense than in villous CTB (Fig. 4D). TEAD4 signal also showed strong localization within proximal anchoring column trophoblasts, while FOXO4 showed a disorganized/diffuse and less intense signal in all aspects of the villous column (Fig. 4D). Consistent with this staining pattern, both FOXO4 and TEAD4 showed strong nuclear signals in CTB-like layers of PD- and hTSC-TOrgs, where FOXO4 showed signal in SCT-like nuclei within the organoid core in both organoid systems (Fig. 4D). Together, these findings confirm that these SCENIC-identified driver TFs are expressed in progenitor CTB-like cells in both PD- and hTSC-TOrgs.

Our finding that cells of the TSC state in hTSC-TOrgs function as upstream progenitors akin to progenitor CTB *in vivo* prompted us to explore trajectory and gene-regulatory relationships between the TSC and CTB states in hTSC-TOrgs. To do this, we subset and re-clustered hTSC-TOrg CTB and TSC and re-assessed splice kinetics using scVelo. scVelo velocity vector length provides an indication of differentiation rate. As well, scVelo enables the identification of putative driver genes that regulate differentiation progress^34^. As expected, the velocity vector field originated in the TSC state and progressed towards the CTB state (Fig. 4E) with an average velocity vector length of 14.48 (Fig. S3C). Interestingly, when the TSC state was clustered with *in vivo* CTB states and PD-TOrg trophoblasts and subjected to scVelo, directionality between these states was not observed, suggesting that the TSC state in PD-TOrgs is not upstream of CTB like it is in hTSC-TOrgs (Fig. S3D). *GCM1* was shown to be a key driver of hTSC-TOrg TSC to CTB state differentiation (Fig. 4F). Comparing expression of CTB-specific TFs in the TSC state, we found comparable TF expression between hTSC-TOrg TSC and MPI/PD-TOrg CTB with the exception of *EGR1* and *TFAP2B* (Fig. 4G). In terms of TSC state maintenance, *TEAD4* and TSC state specific/enriched *SOX13* and *SOX15* TFs were identified (Fig. 4G). Altogether, these findings show that while conservation of key TF drivers exist between *in vivo* and TOrg CTB, differences in regulon rank, level of expression, and hierarchical order of CTB and TSC states in PD- and hTSC-TOrgs highlights possible discrepancies in regulatory features driving model differentiation and maintenance.

### Assessment of transcriptional drivers of the extravillous pathway in TOrgs

EVT differentiation originates from progenitor-like column trophoblasts residing in the proximal column of anchoring villi^21, 35, 36^. Within this niche, EVT progenitors proliferate and undergo an epithelial-to-mesenchymal transition that generates non-proliferative and invasive subsets of EVT. ITGA2 has been proposed as a putative cell surface marker of EVT progenitors in anchoring columns^21^. Because the hTSC-TOrg progenitor TSC state is predominantly *ITGA2^+^*, we compared *in vivo* and TOrg *ITGA2* expression. We found 5% of *in vivo* trophoblasts, mainly those residing in the cCTB state, were *ITGA2^+^,* whereas 20% of PD-(mainly cCTB) and 33% of hTSC-TOrg trophoblasts (spread across all non-SCTp states) were *ITGA2^+^* (Fig. 5A; 5B). We next compared *ITGA2^+^* trophoblast gene signatures between MPI, PD-, and hTSC-TOrg trophoblasts. Overall, some similarity between TOrg and *in vivo* trophoblasts was found with 16,349 common transcripts (Fig. 5C). hTSC-TOrg *ITGA2^+^* gene signatures, however, were the most distinct with 3109 unique transcripts expressed relative to only 334 and 513 unique transcripts in *ITGA2^+^* PD-TOrg and *in vivo* trophoblasts, respectively (Fig. 5C; Table S4D). To verify these findings, we used *in situ* hybridization to probe for *ITGA2* mRNA in chorionic villi as well as PD- and hTSC-TOrg cultures (Fig. 5D). We localized *ITGA2* transcripts to regions of the villous core and clusters of cells in the proximal region of anchoring chorionic villi, although these niches were not found in every column assessed. Compared with this, TOrgs supported our computational findings with overall enrichment of *ITGA2* in CTB-like cells (Fig. 5D). Beyond *ITGA2*, column-associated *TMSB4X* and *EPCAM* transcripts were also upregulated in the TSC state (Fig. S5A).

**Figure 5:**
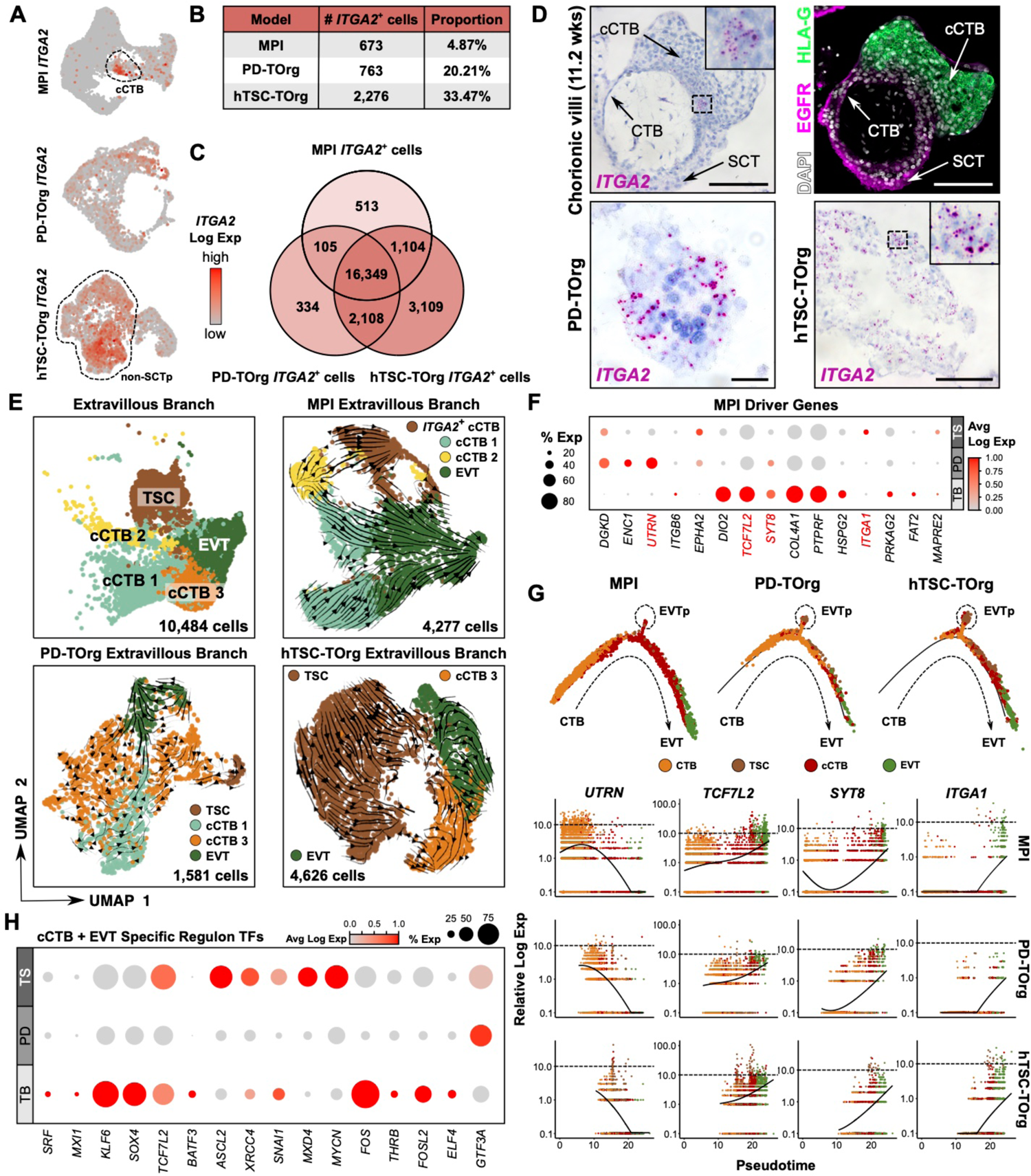
Comparing *in vivo* and TOrg extravillous trophoblast differentiation. **(A)** Log *ITGA2* expression (red) plotted in re-clustered MPI, PD-, and hTSC-TOrg UMAP space. **(B)** Table showing the model-specific number and proportion of *ITGA2^+^* trophoblasts. **(C)** Venn diagram comparing the bulk *ITGA2^+^* MPI, PD-, and hTSC-TOrg transcriptomes. **(D)** Representative *in situ* hybridization images showing *ITGA2* mRNA signal in chorionic villi (GA: 11.2 weeks), PD-(P1111), or hTSC-TOrg (CT29). Within chorionic villus and organoid sections, IF imaging of EGFR and HLA-G are also shown; nuclei are labelled with DAPI. Bar = 100 µm. **(E)** Integrated and re-clustered model-specific extravillous TB UMAP plots. UMAP plots are overlain with RNA velocity vector fields. **(F)** Dot plot comparing the percent and average expression level of genes driving the MPI cCTB and EVT vector fields *in vivo* and in TOrgs. **(G)** Integrated Monocle2 pseudotime ordering of trophoblasts along the extravillous path of differentiation (top) with expression kinetics of select MPI gene drivers plotted across pseudotime in MPI, PD-, and hTSC-TOrgs (bottom). **(H)** Dot plot showing the percent and average expression level of TFs driving cCTB- and EVT-specific regulons *in vivo* and in TOrgs.

We next compared *in vivo* and TOrg cCTB and EVT states. Within the cCTB state, PD- and hTSC-TOrg cCTB-like cells shared 18,193 transcripts with *in vivo* cCTB cells while 264 and 2526 unique transcripts were found in PD- and hTSC-TOrgs, respectively (Fig. S5B; Table S4E). Comparing *in vivo* and TOrg EVT, some differences in gene expression were observed: 15,031 common EVT transcripts were found, and 1629, 270, and 1657 unique transcripts were found in MPI, PD-, and hTSC-TOrg EVT, respectively (Fig. S5C; Table S4F). These differences could result from a lack of non-trophoblast decidual factors *in vitro* that are known to regulate EVT differentiation ^37^. To Assess this, we re-clustered all EVT into 2 states, one state representative of chorionic villous EVT and a second state containing EVT derived from villous and decidual tissues (Fig. S5D). Comparing non-decidual *in vivo*

EVT to TOrg EVT by PCC revealed high transcriptomic similarity: PD-TOrg EVT were 95% similar and hTSC-TOrg EVT were 91% similar to non-decidual *in vivo* EVT (Fig. S5E). Next, we used scVelo to compare *in vivo* and TOrg gene drivers of EVT differentiation. To do this, MPI, PD-TOrg, and hTSC-TOrg cCTBs and EVTs from the integrated object were subset, individually re-clustered, and velocity vector fields were reconstructed (Fig. 5E). Comparing routes of EVT differentiation, *in vivo* trajectory complexities were simplified *in vitro*. However, both *in vivo* and in TOrgs, differentiation progressed through either *ITGA2^+^* cCTB or *ITGA2*^-^ cCTB (Fig. 5E). scVelo-informed drivers of *in vivo* EVT differentiation were then compared with TOrgs. Similar to our exploration of the CTB niche, *in vivo* EVT driver gene expression patterns were conserved *in vitro* with variances presenting at transcript expression level (Fig. 5F). Some differences were observed, however, with overall enrichment of *DGKD*, *ENC1*, *UTRN*, and *EPHA2* transcripts in TOrgs (Fig. 5F).

While scVelo re-assessment allowed for model-specific comparison of EVT differentiation complexities, direct model comparison was not achieved. To address this, we re-constructed Monocle2 extravillous trajectories using CTB, TSC, cCTB, and EVT states derived from all models. This approach unveiled a distinct *EPCAM^+^*, *ITGA2^+^* EVT progenitor cell niche that we termed EVTp (Fig. 5G; S5F). *In vitro*, this EVTp population expands and is predominated by cells of the TSC state, suggesting that the TSC state represents an EVT progenitor-like cell population. Further, we measure an accelerated pseudotime EVT differentiation in hTSC-TOrgs compared to *in vivo* trophoblasts, due to the downstream hTSC-TOrg TSC state origin as well as the general lack of TOrg CTB (Fig. 5G). Using Monocle2 ordering, we compared scVelo-informed *in vivo* gene driver expression across pseudotime and found overall conservation *in vitro*. However, different expression kinetics, due to altered TOrg pseudotime origins, were found (Fig. 5G). Finally, we compared the expression level of TFs driving cCTB- and EVT-specific regulons *in vivo* and in TOrg cCTB and EVT. Overall, *in vivo* TF expression patterns were conserved in TOrgs with the exceptions of *BATF3* found to be specific *in vivo* and *ASCL2* and *MYCN* found enriched in hTSC-TOrgs (Fig. 5H). Together, our results demonstrate high conservation of the non-decidual *in vivo* EVT transcriptome in both TOrg models and highlights differences in cellular hierarchy in EVT development.

### Assessment of transcriptional drivers of the villous pathway in TOrgs

Sub-setting and re-clustering CTB and SCTp states from the integrated trophoblast object resolved two transcriptionally distinct SCTp states (Fig. 6A; Fig. S6A); the SCTp1 state shows conservation *in vivo* and *in vitro*, while the SCTp2 state is predominant in hTSC-TOrgs. Because of this, we compared *in vivo* and TOrg SCTp transcriptomic signatures and routes of villous maturation. Differences in hTSC-TOrg SCTp gene expression were found with 3,459 unique genes expressed in hTSC-TOrg, but only 722 and 126 unique genes expressed *in vivo* and in PD-TOrg SCTp, respectively (Fig. S6B; Table S4G). Considering expression level, higher levels of fusogenic *ERVV-1* and *ERVFRD-1* transcripts were found in the SCTp1 state while higher levels of hormonal *CGA*, *CGB*, *LEP*, and placenta-specific (i.e., *PSG*) transcripts were found in SCTp2 (Fig. S6C), suggesting these unique states represent functional or developmental timing differences during syncytial formation. Comparing *in vivo* and TOrg SCTp1 states by PCC revealed high transcriptomic similarity: PD-TOrg SCTp1 was 93% and hTSC-TOrg SCTp1 was 92% similar to *in vivo* SCTp1 (Fig. S6D).

**Figure 6:**
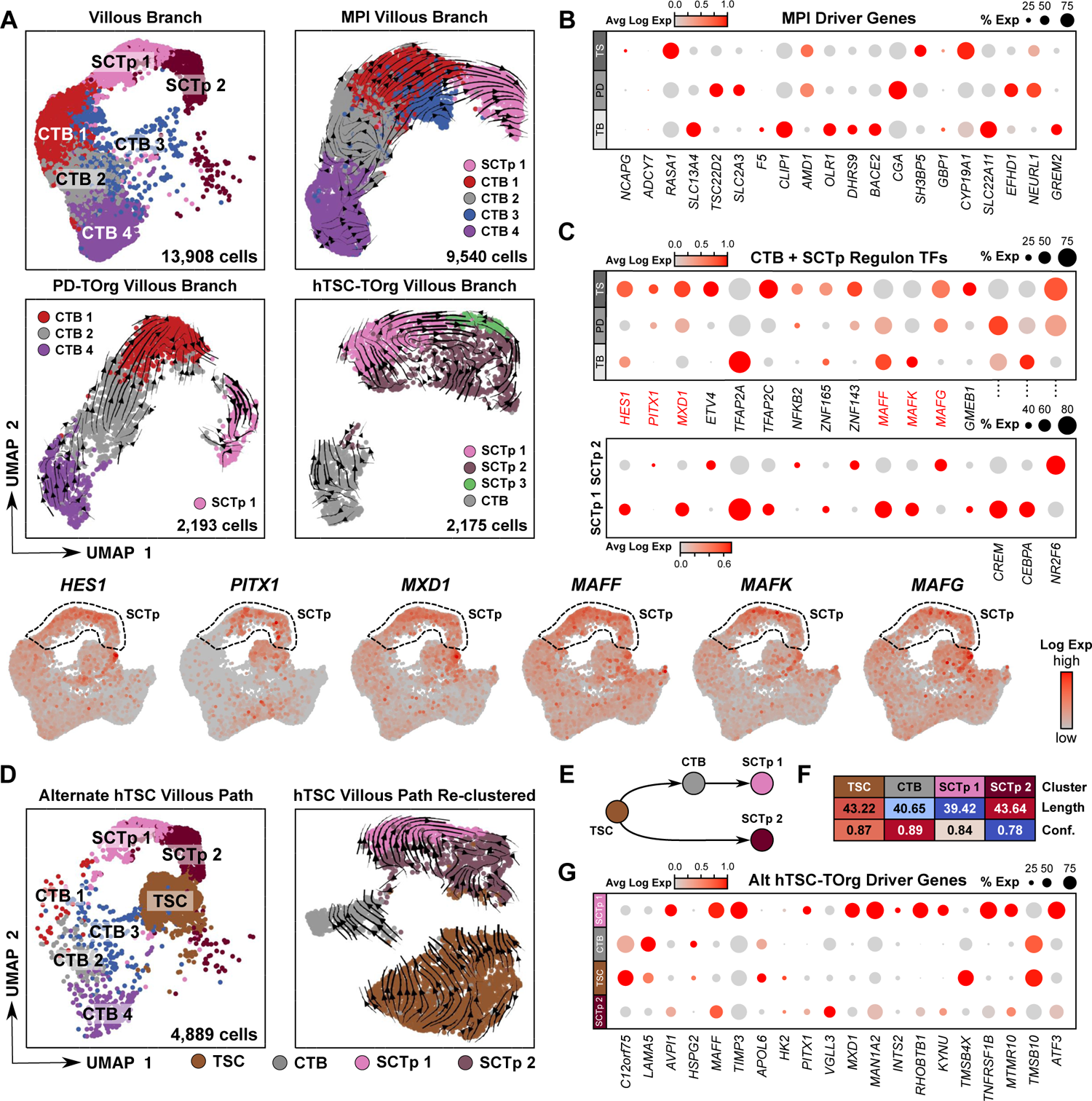
Comparing *in vivo* and TOrg villous trophoblast differentiation. **(A)** Integrated and re-clustered model-specific villous TB UMAP plots. UMAP plots are overlain with RNA velocity vector fields. **(B)** Dot plot comparing the percent expression and average expression level of genes driving the MPI CTB and SCTp vector fields *in vivo* and in TOrgs. **(C)** Dot plot showing the percent and average expression level of TFs driving SCTp-specific regulons in each model. Select TFs are highlighted with expression (red) plotted in integrated UMAP space. **(D)** Integrated CTB, TSC, and SCTp UMAP plot (left), re-clustered (right). **(E)** Novel hTSC-TOrg SCT differentiation paths. **(F)** Heatmap quantifying average state vector length and confidence for the UMAP presented in panel D. **(G)** Dot plot showing the percent and average expression level of genes driving the CTB and SCTp vector fields.

Next, we re-applied scVelo to compare the routes and transcriptional drivers of villous differentiation in TOrgs (Fig. 6A). Overall, PD-TOrgs better recapitulated *in vivo* CTB to SCTp development (Fig. 6A). Increased SCTp heterogeneity, greater distance mapped between CTB and SCTp states in 2D UMAP space, and unconnected CTB and SCTp velocity vectors were observed in hTSC-TOrgs, suggesting that hTSC-TOrg SCT differentiation is more complex than what is found *in vivo* and in PD-TOrgs (Fig. 6A). Gene expression patterns of scVelo-informed *in vivo* drivers of SCTp differentiation were conserved *in vitro* with variances presenting at the level of transcript expression (Fig. 6B). We next compared the expression level of TFs driving SCTp-specific regulons between *in vivo* and TOrg SCTp. Overall, *in vivo* TF expression patterns were conserved in TOrgs with the exception of *PITX1*, *ETV4*, *NFKB2*, *ZNF143*, *MAFG*, and *NR2F6* that were found enriched in both TOrg platforms (Fig. 6C).

Finally, because the TSC state in hTSC-TOrgs appears upstream of the CTB state, we explored how inclusion of the TSC state alters the hTSC-TOrg villous trajectory model. To do this, we subset, re-clustered, and re-constructed velocity vector fields in CTBs, SCTp, and TSCs (Fig. 6D). This resolved two differentiation routes progressing toward mature SCTp, both originating from the TSC state. The first path followed a step-wise progression with TSCs first differentiating into CTB followed by progression into SCTp1 (Fig. 6D; Fig. 6E). The second route showed a direct path originating from TSC to SCTp2 (Fig. 6D; Fig. 6E). Quantified, velocity vectors were longest following this second path (Fig. 6F), suggesting that hTSC-TOrgs prioritize this later route. Notably, a direct TSC state to SCTp2-state pathway was not observed *in vivo* or in PD-TOrg TSCs (Fig. S6E). As well, when *in vivo* and hTSC-TOrg CTB and SCTp states were chimerically re-clustered with hTSC-TOrg TSC and SCTp states, conventional CTB to SCTp differentiation was prioritized (Fig. S6F). We find *TMSB4X* transcripts driving hTSC-TOrg differentiation out of the TSC state (Fig. 6G); *LAMA5*, *HSPG2*, and *TMSB10* transcripts driving differentiation towards CTB; and *TIMP3*, *MXD1*, and *ATF3* driving conventional CTB maturation into SCTp1 (Fig. 6G). Alternatively, *HK2* and *VGLL3* transcripts were identified as drivers of direct TSC to SCTp2 differentiation (Fig. 6G). Overall, these findings highlight that while modest differences exist at the level of gene expression across *in vivo* and *in vitro* syncytium development, PD-TOrgs better recapitulate the cell trajectory order observed *in vivo*. In contrast, while hTSC-TOrgs show a conserved SCTp1 endpoint, a key difference in cell state starting point and an alternative SCTp differentiation pathway highlight limitations of hTSCs in modeling the villous pathway.

## DISCUSSION

By applying a single-cell transcriptomic approach, this study directly and comprehensively compares newly developed human TOrg culture systems with reference to *in vivo* trophoblasts. While TOrgs established from primary first trimester chorionic villous tissue and commercially available hTSC-lines formed self-organizing 3D structures consisting of all trophoblast sub-lineages, cell-state categorization, cell trajectory modeling, and assessment of transcriptional drivers underlying stem cell maintenance and differentiation highlighted notable differences in both primary and hTSC-line organoids. Overall, this work describes conserved and discordant features of current TOrg systems, and in doing so, provides an important resource for researchers needing to interpret published work and evaluate state-of-the-art *in vitro* trophoblast systems in the design of their research. This data is provided as a public resource at: https://humantrophoblastmodels.shinyapps.io/mjs_trophoblasts_2022/.

Previous work comparing 2D and 3D stem cell-based trophoblast cultures identified model-specific differences between primary CTB-derived and hTSC-derived TOrgs^11^. However, because trophoblasts encompass a heterogenous composition of specialized cell subtypes^12, 13^, it was unclear if these differences existed purely at the transcriptome level or were the result of differences in cell frequency or cell type. Our single-cell transcriptomic comparisons revealed that PD-TOrgs better reflect *in vivo* trophoblast composition and differentiation. This observation is in part due to the high prevalence of a distinct cell state in hTSC-TOrgs that is not present in substantial quantities in chorionic villi and PD-TOrgs, which we term the TSC state. Notably, this state concurrently shows progenitor- and column-like transcriptomic features exemplified by high levels of *TEAD4*, *ITGA2*, *NOTCH1*, *EPCAM*, and *TMSB4X*. These features are consistent with EVT progenitors residing in proximal regions of anchoring columns^21, 35, 36^. Like CTB-like progenitors in PD-TOrgs that show lineage trajectory potential for establishing SCTp- and EVT-like states, the TSC state appears to function as a multi-potent progenitor population capable of differentiation into CTB-, SCTp-, and EVT-like states.

We rationalize that TSC state cells are favoured during hTSC clonal expansion and establishment^38, 39^. This selection bias, perhaps the result of cell culture matrix components, pathway inhibitors, and growth factors, favors this proliferative EVT progenitor cell population. Supporting this, we find high levels of proliferation-associated *MKI67* and *CCNA2* transcripts in cells of the TSC state. The negligible presence of the TSC state *in vivo* suggests that TOrg culture conditions may bias EVT progenitor cell gene expression or EVT progenitor cell survival *in vitro*. Consistent with this, TSC-enriched genes (i.e., *ITGA2*, *SOX15*) are localized to the cCTB2-state *in vivo*. cCTB2 clusters near the TSC state when assessed in 2D UMAP space, indicating high transcriptional similarity. Therefore, culture-induced pressures may skew the cCTB2 state towards a TSC-like transcriptome while providing a TOrg fitness advantage that allows for their expansion. Similar selection mechanisms have been observed in intestinal organoids^40^.

Gene regulatory network analyses identified TFs driving key trophoblast progenitor cell regulons. We confirm the importance of known trophoblast progenitor transcripts both *in vivo* and in our TOrgs, including *GATA2/3*^41, 42^, *TFAP2A/C*^41, 42^, *TEAD4*^31–33^, and the newly defined *ARID3A* and *TEAD1*^43^. In addition, our findings highlight novel TFs (i.e., *EGR1*, *TFAP2B*, *FOXP1*, *FOXO4*, *IRF6*) regulating CTB biology *in vivo* and in TOrgs. *FOXO4* provides a fruitful regulatory CTB TF. Interestingly, *FOXO4* has been implicated in redox signalling^44, 45^, tumour suppression^46^, stem cell maintenance^47^, and the inhibition of proliferation through *E2F4*^48^, another CTB-specific TF identified here. Additionally, *FOXO4* regulates rat placenta development^49^ and inhibits epithelial-mesenchymal transition^50^, a process involved in extravillous trophoblast maturation. The regulation of FOXO4 by hypoxia-related processes (i.e., redox signaling, *E2F4* induction) suggests *in utero* hypoxia may induce *FOXO4* to regulate CTB proliferation, renewal, and maintenance, although these interactions will need to be tested.

While PD-TOrgs better recapitulate overall *in vivo* trophoblast development, hTSC-TOrgs appear to best capture extravillous differentiation complexities. The hTSC-TOrg EVT endpoint is transcriptionally similar to the distal column-like CTB or immature EVT observed *in vivo* (i.e., non-decidual EVT). This difference demonstrates the importance of decidual factors in controlling EVT maturation and function^51, 52^, highlighting a limitation in simple TOrg models where signals originating from the maternal environment are difficult to reproduce. Future models incorporating decidual cell (uterine gland epithelia, blood vessel cells, and immune cells) co-cultures may address this limitation, however the integration of such complex models pose significant challenges. Regardless, gene regulatory analyses confirm the importance of *SOX4*, *ASCL2* and *SNAI1* TFs involved in extravillous differentiation *in vivo* and *in vitro*^53–55^ while identifying additional new TF targets (i.e., *TCF7L2*, *BATF3*, *XRCC4*, *MXD4*). Alternatively, computational modeling resolved two routes of TSC maturation towards the SCTp state in hTSC-TOrgs: one conventional CTB to SCTp route defined by fusogenic gene enrichment, and an alternative direct TSC to SCTp route defined by endocrine gene enrichment. Comparing *in vivo* and TOrg regulators of this process, we again confirm known transcripts (i.e., *MSX2*^56^) important for syncytialization while identifying novel (i.e., *PBX1*, *TAF7*, *ESRRA*, *ETV4*) transcripts driving syncytialization *in vivo* and *in vitro*. However, because the placental syncytium represents a large, multinucleated, and diverse cell layer, these *in silico* differences may simply reflect the existence of distinct SCT subtypes found *in vivo*^57, 58^. As well, because TOrg polarity is inverted with an inner SCT layer, TOrg SCT artefacts may bias our single-cell findings. Unfortunately, scRNA-seq and flow cell-sorting limitations prevent the sequencing of true multi-nucleated SCT. Therefore, SCTp similarities and differences reported here might represent only a fraction of the SCT state(s) generated *in vivo* and in TOrgs.

While PD-TOrgs better reflect *in vivo* trophoblast composition and differentiation, there are significant limitations of the PD-TOrg system. Notwithstanding the absence of an intact maternal environment, PD-TOrg establishment relies upon access to first-trimester trophoblast tissue. This access is often limited, and in some cases, ethically constrained. Further, PD-TOrgs, unlike hTSC-TOrgs, are multi-clonal and therefore have greater cell heterogeneity. While this feature is a large advantage for PD-TOrg genetic diversity, it does provide a logistical barrier for studies aiming to implement gene editing strategies like CRISPR-Cas9 that necessitate clonal expansion of genetically edited cells^59^. Additionally, PD-TOrgs are often established from electively terminated first-trimester samples. Therefore, the *in utero* viability and genetic integrity of these samples is not guaranteed^60^. Similar limitations exist for the commercially available hTSC lines^8^. Addressing this, trophoblast stem cell cultures derived from primed embryonic and induced pluripotent stem cells are less susceptible to patient-specific genetic alterations and provide alternative, commercially available models^61, 62^. These models, however, still require careful consideration as they remain susceptible to the same limitations described here in hTSC lines. Nonetheless, both PD- and hTSC-TOrgs arguably provide the best available tools for studying human trophoblast development.

In conclusion, we have identified key differences in trophoblast composition, progenitor trophoblast gene expression, and trophoblast differentiation trajectories between *in vivo* trophoblasts and TOrgs. These differences represent significant factors that need to be carefully considered when selecting an optimal trophoblast model and question the accuracy of using hTSC cultures and hTSC-TOrgs for studying human trophoblast stem cell biology. In making these data accessible, we hope that researchers will be able to use our findings to further optimize current human trophoblast models.

## MATERIALS AND METHODS

### Patient recruitment and tissue collection

Placental tissues were obtained with approval from the Research Ethics Board on the use of human subjects, University of British Columbia (H13-00640). All samples were collected from pregnant individuals (19 to 39 years of age) providing written informed consent undergoing elective terminations of pregnancy at British Columbia’s Women’s Hospital, Vancouver, Canada. First-trimester placental tissues (*n*=15) were collected from participating patients (gestational ages ranging from 5–12 weeks) having confirmed viable pregnancies by ultrasound-measured fetal heartbeat. Patient clinical characteristics i.e., height and weight were obtained to calculate body mass index (BMI: kg/m^2^) and all consenting patients provided self-reported information via questionnaire to having first-hand exposure to cigarette smoke or taken anti-inflammation or anti-hypertensive medications during pregnancy. Patient and sample phenotype metadata are summarized in Table S2A.

### Trophoblast organoid establishment and culture

#### Patient-derived trophoblast organoids

PD-TOrg cultures (*n*=5) were established following a modification of previously described protocols (Haider *et al.*, 2018; Turco *et al.*, 2018). In short, primary chorionic villi were washed using 1X HBSS (Gibco) and gently scraped from the chorionic membrane before 8 minute digestion in 1X HBSS, 1mg/ml DNAse I (Sigma-Aldrich), 0.25% Trypsin-EDTA (Gibco) at 37°C. Following incubation, the supernatant was removed and chorionic villi were digested for 15 minutes at 37°C an additional two times. The supernatant from the second and third digestions were then pooled and strained through a 100 μm cell strainer (VWR). Pooled and digested villous suspensions were then centrifuged at 4°C at a speed of 1200 rpm for 10 minutes. The resulting pellet was resuspended in 5mL of DMEM/F12 (Gibco), 1% Penicillin-Streptomycin (Gibco), 1% Antibiotic-Antimycotic (Gibco), and 10% Fetal Bovine Serum (Gibco) and progenitor cytotrophoblasts were purified by Percoll density gradient centrifugation at 22°C using a speed of 2,350 rpm for 24 minutes. The trophoblast enriched layer was collected and washed twice with 1X HBSS. Trophoblasts were resuspended in undiluted ice-cold growth factor-reduced Matrigel (Corning) and cell suspensions (40 μl) containing 30-100 x 10^3^ cells were seeded in each well of a 24-well plate. Plates were placed at 37°C for 2 minutes. Following this, plates were turned upside down to promote equal spreading of cytotrophoblasts throughout the Matrigel domes. After 15 minutes, plates were flipped (right-side up) and 0.5 ml pre-warmed trophoblast organoid media containing advanced DMEM/F12 (Gibco), 10 mM HEPES, 1 X B27 (Gibco), 1 X N2 (Gibco), 2 mM L-glutamine (Gibco), 100 μg/ml Primocin (Invivogen), 100 ng/ml R-spondin (PeproTech), 0.5 μM A83-01 (Tocris), 100 ng/ml rhEGF (PeproTech), 3 μM CHIR99021 (Tocris) and 1 μM Y-27632 (EMD Millipore) was added to each well. This media is referred to as regenerative Wnt^+^ media. Media was replenished every 3-5 days and cultures were passaged on 14- to 21-day intervals, depending on organoid size and organoid density within the Matrigel dome.

#### Trophoblast stem cell-derived organoids

The hTSC lines CT27 (RCB4936), CT29 (RCB4937), and CT30 (RCB4938) obtained from the Riken BRC Cell Bank were established as 2D cultures in hTSC regenerative Wnt^+^ media^8^. Spontaneously self-organizing trophoblast organoids were established using identical regenerative Wnt^+^ and EVT-differentiating Wnt^-^ media as described for PD-TOrg establishment and differentiation. All hTSC lines were initiated at passage 17 (P17) and did not exceed P24.

#### Trophoblast organoid EVT Differentiation

EVT differentiation in PD-TOrg (*n*=5) and hTSC-TOrg (CT27, CT29, CT30) cultures was performed as described in Shannon et al., 2022. Briefly, organoids were cultured until >50% colonies reached at least 100 µm in diameter (∼5 days for hTSC-TOrgs and ∼10 days for PD-TOrgs) or 200 μm for the organoids used in scRNA-seq workflow. Following this, organoids were maintained in EVT-differentiation Wnt^-^ media containing advanced DMEM/F12, 0.1 mM 2-mercaptoethanol (Gibco), 0.5% penicillin/streptomycin (Gibco), 0.3% Bovine Serum Albumin (Sigma Aldrich), 1% ITS-X supplement (Gibco), 100 ng/mL NRG1 (Cell Signaling Technology), 5.0 µM A83-01 (Tocris), and 4% Knockout Serum Replacement (Gibco). Media was exchanged for EVT-differentiation Wnt^-^ media lacking NRG1 on day 6 of differentiation following visible trophoblast outgrowth. PD- and hTSC-TOrg differentiation was stopped on day 7 of differentiation for 10x single-cell RNA sequencing workflow and flow cytometry analyses. Differentiation was stopped on day 10 of differentiation for all EVT-differentiated organoid qPCR analyses.

### Flow cytometry analysis

Single cells from PD-TOrgs (P1069 and P1111) and hTSC-TOrgs (CT27, CT29, and CT30) were isolated by incubation with Cell Recovery Solution (Corning) for 40 minutes at 4°C. Cells were then washed in ice-cold PBS and dissociated into single cells using TrypLE^TM^ Express (Gibco) supplemented with 5% DNAse I for 7 minutes at 37°C. Following this, organoid cells were filtered and stained with Fix Viability Dye 780 (eBiosciences) for 20 minutes at room temperature for live/dead exclusion. Then, cells were washed in PBS and incubated with Fc Receptor Binding Antibody Inhibitor (1:20; eBiosciences) for 5 minutes at 4°C, followed by 30 minutes incubation with anti-EGFR BV605 (clone AY13; 1:50; Biolegend) and anti-HLA-G APC (clone MEMG-9; 1:150; Abcam). Cells were then washed in PBS and resuspended in FACS buffer (PBS supplemented with 2% FBS and 0.5 mM EDTA). Data was acquired using a Becton Dickinson (BD) FACSymphony A5 and data was analyzed using FlowJo v10.8.1.

### TOrg growth assay

hTSC-TOrgs (CT27, CT29, CT30) and PD-TOrgs (P1069, P1111) were cultured in regenerative Wnt^+^ media as described above. Bright-field images were taken every 3 days on an AxioObserver inverted microscope (Carl Zeiss) using a 5X objective until organoid cultures reached a terminal ∼200 μm diameter (∼9 days for hTSC-TOrgs and ∼15 days for PD-TOrgs). The tiling feature in the Zeiss software (version 3.1) was used to image and measure consistent groups of organoids over the course of the experiment. Four positions (localized using x, y, and z plane coordinates) were selected for each experiment, allowing the same organoids to be followed across culture. Each position contained *n*=5-35 organoids. Organoid size (μm^2^) was measured for all organoids imaged in each position using ImageJ and growth curves were calculated. hTSC-TOrg and PD-TOrg mean areas (μm^2^) for each culture timepoint are summarized in Table S1.

### *ELF5* promoter Pyrosequencing

Bisulfite-pyrosequencing was used to measure the methylation status of 6 different CpGs within the promoter/5’ UTR of *ELF5* that showed differential methylation between human trophoblasts and other placental cell types (i.e., stromal fibroblasts, endothelium, Hofbauer cells) based on our previously published cell-specific characterization of the placental methylome^63^, available online at: https://robinsonlab.shinyapps.io/Placental_Methylome_Browser/.

The bisulfite-pyrosequencing assay was designed using PyroMark Assay Design software version 2.0 with the following primers: *ELF5*-F: 5’-/5Biosg**/**GGTTTTAGGTAGAGTTTTTGAGTTTTGATA-3’; *ELF5*-R: 5’-ACAAAACTACAAATATCTTTATTTCCACT-3’; *ELF5*-Seq: 5’-CTACAAATATCTTTATTTCCACT-3’. The assay was validated using a standard curve calibrated against human methylation controls (EpigenDx, USA Cat. 80-8061-HGHM-5 and 80-8062-HGUM-5) and isolated placental and somatic tissues. Approximately 200-300 ng genomic DNA were bisulfite converted using the EZ DNA Methylation-Gold Kit (Zymo, USA Cat D5006) following manufacturer’s protocol for all steps. Bisulfite-converted genomic DNA was then amplified using HotStar Taq and the PCR product was pyrosequenced on a PyroMark MD instrument (Qiagen, USA). The methylation status of each CpG of interest (*n*=6 per sample) was evaluated with PyroMark Q-CpG software (Qiagen) and data from all CpG sites was averaged for each sample and presented as percent methylation (Figure S1E).

### PD-TOrg single-cell RNA-seq library construction

For all organoid cultures, media was aspirated and patient-derived trophoblast organoids were diluted in Cell Recovery Solution at 4°C for 40 minutes on either culture day 7 in regenerative Wnt^+^ media (for non-differentiated organoids) or culture day 7 in EVT-differentiation Wnt^-^ media (for EVT-differentiated organoids) after initial establishment to 200 μm in regenerative Wnt^+^ media. Cells were then washed with cold PBS and centrifuged at 1000 rpm for 5 minutes. Cells were disaggregated in TrypLE containing 5% DNase I at 37°C for 7 minutes before the addition of either 100 μl regenerative Wnt^+^ media (for non-differentiated organoids) or 100 μl EVT-differentiating Wnt^-^ media (for EVT-differentiated organoids). The resulting cell suspensions were then sieved through a 40 μm filter before library construction.

#### Library construction and sequencing

Single-cell suspensions were stained with 7-Aminoactinomycin D (7-AAD) viability marker and live/dead sorted using a Becton Dickinson (BD) FACS Aria. Cell suspensions were loaded onto the 10x Genomics Chromium Controller for GEM bead generation and single-cell barcoding. Non-differentiated and differentiated primary cytotrophoblast-derived organoid single-cell libraries were prepared using the 10x Chromium Next GEM single-cell 3’ reagent chemistry kit v3.1, following manufacturer’s protocol for all steps. Single-cell libraries were sequenced on an Illumina Nextseq 500 instrument at a sequencing depth of 150,000 reads/cell. For all samples, the 10X Genomics Cell Ranger version 3.0 software was used for STAR read alignment as well as barcode and UMI counting against the hg19 reference genome.

#### Data Repository Integration

Droplet-based first-trimester scRNA-seq decidual and placental tissue data was obtained from the public Gene Expression Omnibus (GEO) repository (GEO174481) and from the ArrayExpress public repository (E-MTAB-6701)^12, 13^. Raw BAM files from the GEO and ArrayExpress repositories were downloaded and pre-processed using 10X Genomics Cell Ranger version 3.0 using identical parameters as described in Shannon et al., 2022, again against the hg19 reference genome. Sample specific sequencing metrics are summarized in Table S2B.

### Single-cell RNA-seq data analysis

The code used for all subsequent data processing and analyses are available at: https://github.com/MatthewJShannon.

#### Data pre-processing and quality control

3774 cells from PD-TOrgs (3,049 cells from undifferentiated TOrgs and 725 cells from EVT-differentiated TOrgs) were sequenced and pre-processed using the Seurat R package (version 4.0.1)^64, 65^ and integrated with publicly available raw data (available at: GEO174481 and E-MTAB-6701) published in Shannon et al., 2022. Additional chorionic villi EPCAM^+^ CTBs and placental and decidual HLA-G^+^ EVTs that were not included in Shannon et al., 2022, were accessed from E-MTAB-6701 and included here. To ensure only high-quality cells were used in downstream analyses, cells containing fewer than 500 detected genes and greater than 20% mitochondrial DNA content were removed. Individual samples were scored based on expression of G_2_/M and S phase cell cycle gene markers, scaled, normalized, and contaminating cell doublets were removed using the DoubletFinder package version 2.0.3^66^ with default parameters and a doublet formation rate estimated from the number of captured cells. To minimize bias due to read dropouts the “FindVariableFeatures” function was used with a “vst” selection method to prioritize highly variable genes in each sample for all downstream analyses. Following pre-processing, all samples were merged and integrated using cell pairwise correspondences between single cells across sequencing batches. During integration, the Pearson residuals obtained from the default regularized negative binomial model were re-computed and scaled to remove latent variation in percent mitochondrial DNA content as well as latent variation resulting from the difference between the G_2_/M and S phase cell cycle scores.

#### Trophoblast sub-setting, cell clustering, and identification

1. *Integrated trophoblast object:* Trophoblasts were identified if they fulfilled one of two criteria: First, trophoblasts were selected if they expressed *KRT7, EGFR, TEAD4, TP63, TFAP2A, TFAP2C, GATA2, GATA3, HLA-G, ERVFRD-1* transcripts at a level greater than zero, and *VIM, WARS, PTPRC, DCN, CD34, CD14, CD86, CD163, NKG7, KLRD1, HLA-DPA1, HLA-DPB1, HLA-DRA, HLA-DRB1, HLA-DRB5, HLA-DQA1* transcripts at a level equal to zero. Second, trophoblasts were selected if they originated from either the PD-TOrg or hTSC-TOrg *in vitro* trophoblast models, regardless of gene expression. After identification, trophoblasts were subset and re-clustered in Seurat at a resolution of 0.375 using 30 principal components. Cell identity was determined through observation of known trophoblast gene markers (*CGB*, *ERVFRD-1*, *HLA-G*, *ITGA5*, *EGFR*, *TP63*, *ELF5*, *BCAM*, etc.) as well as through observation of proliferative gene markers (*CCNA2* and *MKI67*) in each cluster and within each model (Fig. 1G). Cell state identity was further confirmed using cluster marker genes found via the “FindAllMarkers” function using a model-based analysis of single-cell transcriptomics (MAST) GLM-framework implemented in Seurat^67^. Gene markers were selected as genes with a minimum log_2_ fold change value > 0.25 and an adjusted p-value < 0.05 within a specific cluster. Identified cluster marker genes are summarized in Table S3A for the integrated trophoblast object. Trophoblast projections were visualized using Uniform Manifold Approximation and Projections (UMAPs).
2. *Maternal-placental interface (in vivo) trophoblast object:* Primary placenta (chorionic villi) and decidua-derived trophoblasts were subset from the integrated trophoblast object and re-clustered in Seurat at a resolution of 0.375 using 30 principal components. Similar to the integrated trophoblast object, *in vivo* trophoblast cluster identity was determined through observation of known trophoblast gene marker expression, proliferative gene marker expression, and from the unique cluster gene markers identified through MAST differential expression analysis in Seurat, as described. Identified cluster marker genes are summarized in Table S3B. For all *in vivo* trophoblast analyses (i.e., ligand-receptor analyses, pseudotime, etc.), identified *in vivo* states were binned as follows: CTB states 1-4 were merged into one CTB state and cCTB states 1 and 2 were merged into one cCTB state. This created 4 states representative of each *in vivo* trophoblast-subtype (CTB, SCTp, cCTB, and EVT). Trophoblast projections were visualized using UMAPs.
3. *PD-TOrg trophoblast object:* PD-TOrg-derived trophoblasts were subset from the integrated trophoblast object and re-clustered in Seurat at a resolution of 0.150 using 30 principal components. Similar to the integrated trophoblast object, PD-TOrg trophoblast cluster identity was determined through observation of known trophoblast gene marker expression, proliferative gene marker expression, and from the unique cluster gene markers identified through MAST differential expression analysis in Seurat, as described. Identified cluster marker genes are summarized in Table S3C. For all PD-TOrg analyses (i.e., ligand-receptor analyses, pseudotime, etc.), identified PD-TOrg states were binned as follows: CTB states 1 and 2 were merged into one CTB state. This created 5 states representative of each PD-TOrg trophoblast-subtype (CTB, TSC, cCTB, EVT, and SCTp). Trophoblast projections were visualized using UMAPs.
4. *hTSC-TOrg trophoblast object:* hTSC-TOrg-derived trophoblasts were subset from the integrated trophoblast object and re-clustered in Seurat at a resolution of 0.250 using 30 principal components. Similar to the integrated trophoblast object, hTSC-TOrg trophoblast cluster identity was determined through observation of known trophoblast gene marker expression, proliferative gene marker expression, and from the unique cluster gene markers identified through MAST differential expression analysis in Seurat, as described. Identified cluster marker genes are summarized in Table S3D. For all hTSC-TOrg analyses (i.e., ligand-receptor analyses, pseudotime, etc.), identified hTSC-TOrg states were binned as follows: TSC states 1 and 2 were merged into one TSC state and SCTp states 1 and 2 were merged into one SCTp state. This created 5 states representative of each hTSC-TOrg trophoblast-subtype (CTB, TSC, cCTB, EVT, and SCTp). Trophoblast projections were visualized using UMAPs.
5. *CTB trophoblast subset:* CTB states 1-4 were subset from the integrated trophoblast object and subsequently subset based on model, i.e.: Maternal-Placental interface (*in vivo*) CTBs (8,465 cells), PD-TOrg CTBs (1,921 cells), or hTSC-TOrg CTBs (598 cells). *In vivo* and PD-TOrg CTBs were each re-clustered in Seurat at a resolution of 0.250 using 30 principal components and hTSC-TOrg CTBs were re-clustered in Seurat at a resolution of 0.150 using 30 principal components. Cell state identity was determined by comparing expression of the CTB and proliferative gene markers used to identify each CTB state in the integrated object. Next, CTB states 1-4 as well as the TSC state were subset from the integrated trophoblast object and further subset by model into either *in vivo* progenitor cells (8,505 cells), PD-TOrg progenitor cells (2,051 cells), or hTSC-TOrg progenitor cells (3,312 cells), for assessment of TSC’s influence on trophoblast progenitor CTB. *In vivo* and PD-TOrg progenitors were each re-clustered in Seurat at a resolution of 0.250 using 30 principal components while hTSC-TOrg progenitors were re-clustered in Seurat at a resolution of 0.100 using 30 principal components. Cell state identity was similarly determined by comparing expression of the CTB, TSC, and proliferative gene markers used to identify each CTB and TSC state in the integrated object. Model-specific CTB and model-specific CTB with TSC projections were visualized using UMAPs.
6. *Model specific origin subsets:* Cells overlapping trophoblast states expressing the greatest progenitor-associated gene signature *in vivo* (CTB; 4,179 cells), PD-TOrgs (CTB; 1,303 cells), and hTSC-TOrgs (TSC; 2,191 cells) were subset using Seurat following the pipeline summarized in Fig. S2A and used for origin analyses presented in Fig. 3.
7. *Extravillous differentiation branch subset:* cCTB states 1-3 as well as the TSC and EVT states were subset from the integrated trophoblast object and further subset by model into either the Maternal-placental interface (*in vivo*) extravillous branch (4,277 cells), the PD-TOrg extravillous branch (1,581 cells), or the hTSC-TOrg extravillous branch (4,626 cells). Cells of the *in vivo* and PD-TOrg extravillous branches were each re-clustered in Seurat at a resolution of 0.100 using 30 principal components and cells of the hTSC-TOrg extravillous branch was re-clustered in Seurat at a resolution of 0.200 using 30 principal components. Cell state identity was determined in each object by comparing expression of the TSC, cCTB and EVT gene markers used to identify these states in the integrated object. All extravillous differentiation branch projections were visualized using UMAPs.
8. *Villous differentiation branch subset:* CTB states 1-4 as well as SCTp states 1 and 2 were subset from the integrated trophoblast object and further subset by model into either the Maternal-placental interface (*in vivo*) villous branch (9,540 cells), the PD-TOrg villous branch (2,193 cells), or the hTSC-TOrg villous branch (2,175 cells). The *in vivo* villous branch was re-clustered in Seurat at a resolution of 0.250 using 30 principal components, the PD-TOrg villous branch was re-clustered in Seurat at a resolution of 0.100 using 30 principal components, and the hTSC-TOrg villous branch was re-clustered in Seurat at a resolution of 0.200 using 30 principal components. Cell state identity for each object was determined by comparing expression of the CTB and SCT gene markers used to identify these states in the integrated object. Next, CTB1-4, SCTp1-2, and the TSC states were subset from the integrated trophoblast object and further subset by model into either *in vivo* villous (alternative) cells (9,580 cells), PD-TOrg villous (alternative) cells (2,323 cells), or hTSC-TOrg villous (alternative) cells (4,889 cells) to assess the influence of the TSC state on trophoblast villous differentiation in each model. The alternative *in vivo* villous branch was re-clustered in Seurat at a resolution of 0.250 using 30 principal components, the alternative PD-TOrg villous branch was re-clustered in Seurat at a resolution of 0.100 using 30 principal components, and the alternative hTSC-TOrg villous branch was re-clustered in Seurat at a resolution of 0.150 using 30 principal components. Cell state identity was similarly determined by comparing expression of the CTB, SCT, and TSC gene markers used to identify these states in the integrated object. Finally, the *in vivo* and hTSC-TOrg CTB1-4, SCTp1-2, and TSC states (7,234 cells) were subset from the integrated dataset and chimerically merged to assess if the hTSC-TOrg TSC state can differentiate towards *in vivo* CTB and SCTp states *in silico*. Cells were re-clustered in Seurat at a resolution of 0.450 using 30 principal components. As described, cell state identity was determined by comparing expression of the CTB, SCT, and TSC gene markers used to identify these states in the integrated object. All villous, as well as villous and TSC, combinations were visualized using UMAPs.

#### Pseudo-bulk Data comparison

1. *Model comparison:* Trophoblast expression matrices were independently averaged by model (MPI, PD-TOrg, or hTSC-TOrg) and log transformed to create model-specific “pseudo-bulk” gene expression data. Pearson correlation coefficients of the relationship between these pseudo-bulk samples was calculated using the “cor” function from the stats R package (version 4.0.5).
2. *Integrated trophoblast cell type comparison:* Trophoblast expression matrices were independently averaged by integrated trophoblast-subtype (CTB, TSC, cCTB, EVT, or SCTp) and log transformed to create trophoblast state-specific “pseudo-bulk” gene expression data. Pearson correlation coefficients of the relationship between these pseudo-bulk trophoblast subtypes was calculated as described.
3. *Model-specific origin comparison:* Model-specific trophoblast origins were identified and subset as described in Fig. S2B. Expression matrices from these origins were then independently averaged and log transformed to create “pseudo-bulk” MPI, PD-TOrg, or hTSC-TOrg origin gene expression data. Pearson correlation coefficients quantifying the gene expression similarity between these pseudo-bulk origins was calculated as described.
4. *Non-decidual EVT comparison:* Expression matrix data from the integrated, re-clustered, EVT1 state was independently averaged by trophoblast model and log transformed to create model-specific non-decidual EVT “pseudo-bulk” gene expression data. Pearson correlation coefficients of the relationship between pseudo-bulk EVT was calculated as described.
5. *Fusogenic SCTp comparison:* Expression matrix data from the SCTp1 state was independently averaged by trophoblast model and log transformed to create model-specific SCTp1 “pseudo-bulk” gene expression data. Pearson correlation coefficients of the relationship between pseudo-bulk SCTp1 was calculated as described.

#### Venn-Diagrams

To compare pseudo-bulk and cell-specific transcriptomic signatures between each trophoblast model, trophoblasts from each model (i.e., MPI, PD- or hTSC-TOrg) were subset. Next, average gene expression values were calculated for every gene in every trophoblast selected in each condition. From this list, genes expressed at a level greater than 0 were selected as expressed. The VennDiagram package (version 1.7.1) was then used to generate a Venn-diagram showing genes shared, or distinct to, each model based on binarized expression independent of expression level. Venn-diagrams were exported as .svg files for figure generation. The “get.venn.partitions” function was used to extract the list of gene names found in each Venn-diagram quadrant and these lists were exported from R as excel files for table generation. Quadrant-specific gene names for each Venn-diagram are summarized in Table S4.

#### Differential gene expression

Differential expression (DGE) analyses were performed in Seurat using the “FindMarkers” function using a MAST framework generated on the expression matrix data. To include the whole cell transcriptome in DGE analysis, parameters were set to include: genes with a log_2_ fold-change difference between groups of “−INF”, genes with a minimum gene detection of “−INF” in all cells in each group, and genes with a minimum difference in the fraction of detected genes within each group of “−INF”. Results were visualized using the R package EnhancedVolcano (version 1.8.0) with a fold-change cut-off of 1.00 and a p-value cut off of 10e-50 taken as the thresholds for significance for the DGEs presented in Fig. 3E. Comprehensive DGE results are summarized in Table S6.

#### Ligand-Receptor Interactions

The Connectome^16^ version 1.0.1 R package was used to assess cell signalling topologies and ligand receptor interactions within first-trimester chorionic villi trophoblasts and in both the PD-TOrg and hTSC-TOrg models. To accomplish this, ligand and receptor genes in each model were identified, mapped, and the “CreateConnectome” function was used to generate a cell signalling connectome for each model. Following this, binned trophoblast-subtype (i.e., CTB, TSC, cCTB, EVT, or SCTp) production and reception of selected gene family signals (ADAM, BMP, EGF, Laminin, Assorted matrix, matrix glycoprotein, MMP, NOTCH, VEGF, and WNT) were assessed using the “Centrality” function. Centrality assessment ranked the incoming and outgoing cell type edge-weights and assessed the associated Kleinberg directional centrality to identify top ligand producers and correlated receptors.

#### Pseudotime Analysis

The Monocle2^25, 26^ R package version 2.18.0 and Monocle3^27^ R package version 1.1.0 were used to explore the differentiation of progenitor trophoblasts into specialized SCTp and EVT endpoints *in vivo* and in our *in vitro* PD-TOrg and hTSC-TOrg models.

1. *Maternal-placental interface (in vivo) trophoblasts:* The *in vivo* trophoblast expression matrices were loaded into Monocle2 for semi-supervised single-cell ordering in pseudotime using *CDX2*, *SPINT1*, *ELF5*, and *BCAM* expression to denote cells at the origin as well as *ERVFRD-1*, *GCM1*, and *SDC1* gene expression to indicate progress towards SCT, and *HLA-G*, *GCM1*, and *SDC1* gene expression to indicate progress towards EVT. Branched expression analysis modelling (BEAM) was applied to observe the expression of known trophoblast-subtype gene markers (CTB: *BCAM*, *SPINT1*, *TP63*, *ELF5*, *TEAD4*; SCT: *ERVW-1*, *ERVFRD-1*, *CGB*; cCTB: *NOTCH1*, *ITGA2*, and; EVT: *NOTCH2*, *ITGA5*, *HLA-G*, *ITGA1*) across pseudotime, progressing towards either mature EVT or SCTp. Results were visualized using the “plot_cell_trajectory” function. To resolve trophoblast differentiation trajectories in each model at higher resolution, we applied the Monocle3 package version 1.1.0 (Cao *et al.*, 2019). Trophoblast expression matrices were input, clustered, and the principal graph was created using the “learn_graph” function. The “order_cells” function was then used to order cells in pseudotime, designating the trophoblast CTB root based on results from Monocle2 pseudotime ordering and the progenitor gene expression signatures mapped in Fig. S2A.
2. *PD-TOrg trophoblasts:* PD-TOrg pseudotime ordering was conducted as described above for both Monocle2 and Monocle3.
3. *hTSC-TOrg trophoblasts:* The hTSC-TOrg trophoblast expression matrices were loaded into Monocle2 for semi-supervised single-cell ordering in pseudotime using *SPINT1*, *ELF5*, and *BCAM* expression to denote cells at the origin as well as *ERVFRD-1*, *GCM1*, and *SDC1* gene expression to indicate progress towards SCT, and *HLA-G*, *GCM1*, and *SDC1* gene expression to indicate progress towards EVT. *CDX2* was not used here due to low hTSC-TOrg expression that resulted in *CDX2* being filtered out of the dataset during initial variable feature selection prior to pseudotime ordering. Branched expression analysis modelling (BEAM) was applied as described to observe the expression of known trophoblast-subtype gene markers across pseudotime, progressing towards either mature EVT or SCTp. Results were visualized as described. Monocle3 was applied as described, designating the trophoblast root within the TSC state based on results from Monocle2 pseudotime ordering and the progenitor gene expression signatures mapped in Fig S2A.
4. *Integrated extravillous branch trophoblasts:* The integrated CTB, TSC, cCTB, and EVT trophoblast state raw cell counts were loaded into Monocle2 for semi-supervised single-cell ordering in pseudotime using *SPINT1*, *ELF5*, and *BCAM* expression to denote cells at the origin and *HLA-G*, *GCM1*, *SDC1*, and *ITGA1* gene expression indicating progress towards mature EVT. Direct comparison of identified *in vivo* EVT differentiation gene driver expression across pseudotime between *in vivo* and TOrg trophoblasts was achieved using the “plot_genes_in_pseudotime” function, splitting the plot by model (MPI, PD-, or hTSC-TOrg).

#### Lineage Trajectory Analysis

The Velocyto^34^ package was run in terminal using the command line interface to generate sample-specific files containing count matrices of unspliced, spliced, and ambiguous RNA transcript abundances. The velocyto.R (version 0.6.0) implementation of the package was then used to merge each file with its corresponding 10X sample during data pre-processing in Seurat, described above. The final, processed data object and UMAP embeddings were exported to a Jupyter Notebook and the scVelo^68^ Python package was used to read, prepare, and recover holistic trophoblast gene splicing kinetics. Velocity vector fields were projected on top of the extracted UMAP embeddings, generated in Seurat, for each trajectory branch in MPI, PD-, and hTSC-TOrg trophoblast models. The “scv.tl.velocity_confidence” function was used to quantify velocity vector lengths and percent vector confidence for each condition in each model. The “scv.tl.rank_velocity_genes” function was used to identify genes with cluster-specific differential velocity expression, identifying putative gene drivers of progress through the velocity vector field.

#### Regulon enrichment analysis

The SCENIC^29^ (version 1.3.1), AUCell (version 1.18.1), GENIE3 (version 1.18.0), and RcisTarget (version 1.16.0) R packages were used to establish cytotrophoblast-specific gene regulatory networks (GRNs) for MPI, PD-TOrg, and hTSC-TOrg CTBs as well as trophoblast state-specific GRNs for all states identified in the integrated trophoblast object. *cis*-regulatory analysis was conducted on each model and motif enrichment occurred within gene promoters using databases that scored either 500bp upstream of the transcription start site or 20kbp surrounding the transcription start site (±10kbp). Default parameters were used for each SCENIC analysis for each condition. Because SCENIC utilizes a Random Forest approach, results from each run vary^29^. To address this, we ran SCENIC multiple times per condition and present here the aggregated results from all runs. For CTB-specific analyses, we ran SCENIC *n*=5 times each for all PD- and hTSC-TOrg CTBs. Due to computational limitations, the Maternal-placental interface CTB dataset (representing 8,465 total cells) was down-sampled using the “subset” function in Seurat to include a random selection of 500 cells from each CTB state. This provided 2,000 unique *in vivo* CTBs for each SCENIC run (*n*=5). In brief, the GENIE3 package was first used to identify regulatory TFs using the “runGenie3” function. Following this, the “runSCENIC_1_coexNetwork2modules” function was used to generate TF modules, the “runSCENIC_2_createRegulons” function was used to identify genes within each regulon module, and the “runSCENIC_3_scoreCells” function was used to score network activity in each model. To aggregate the CTB-specific results, we combined the ranked TF outputs from each SCENIC run into one table for each model. To reward consistent TF regulons in ranking, a score of 300 was assigned to TFs that were not identified in a respective run, effectively replacing NA ranks. TF ranks were then summed across all 5 SCENIC assessments and scores were ordered in ascending rank to generate final CTB regulon rankings. A confidence score was next calculated for each regulon, representing the relative frequency distribution of each regulon across all 5 SCENIC assessments. Genes regulated in each regulon in the model-specific CTB datasets were determined for each SCENIC analysis. Regulon genes from one representative SCENIC assessment are summarized in Table S7A-C for the CTB-specific analyses.

For the trophoblast state-specific analyses, a similar approach was taken. This time, the integrated trophoblast dataset was down-sampled using the “subset” function in Seurat to include a random selection of 500 cells from each trophoblast state. This provided 5,500 unique trophoblasts for each SCENIC run (*n*=5). To aggregate results and determine the trophoblast-subtype-specificity of each regulon, the scaled regulon activity in each integrated trophoblast state was plotted as a heatmap for each SCENIC run (one representative heatmap is plotted in Fig. S4A). For each regulon, activity in each binned trophoblast-subtype (i.e., CTB, TSC, cCTB, EVT, or SCTp) was determined. Regulon specificity to trophoblast subtypes were compared across all 5 SCENIC runs and regulons were determined as active in a trophoblast-subtype if found active in that subtype in at least 3/5 SCENIC runs. To cross reference these results, the “runSCENIC_4_aucell_binarize” function was used to binarize regulon activity as on/off in each trophoblast state. Binarized percent regulon activity of significant regulons was then plotted as a heatmap for all 5 SCENIC runs with one representative heatmap shown in Fig. S4B. Similar to the CTB-specific analyses, a confidence score was then calculated for each regulon, representing the relative frequency distribution of each regulon across all 5 SCENIC assessments. Trophoblast state-specific results are summarized in Table S7D. Genes regulated in each regulon in the integrated trophoblast dataset were determined for each SCENIC analysis. Regulon genes from one representative SCENIC assessment are additionally summarized in Table S7D for the trophoblast state-specific analyses.

#### Human Trophoblast Model App Generation

The ShinyCell^69^ (version 2.1.0) R package was used to assemble the “human trophoblast models” app and the rsconnect (version 0.8.27) R package was used to publish the app on the shinyapps.io server for public access. The app is available at: https://humantrophoblastmodels.shinyapps.io/mjs_trophoblasts_2022/

### RNA isolation

#### Trophoblast organoids

Total RNA was prepared from PD-TOrg and hTSC-TOrg cultures using TRIzol reagent (Invitrogen) followed by RNeasy Mini kit (Qiagen) clean-up following manufacturer’s protocol. RNA purity was confirmed using a NanoDrop Spectrophotometer (Thermo Fisher Scientific).

#### Bulk chorionic villous tissue

Prior to RNA extraction, human placental chorionic villi were snap frozen in liquid nitrogen. The snap-frozen tissue was then bio-pulverized under super-cooled conditions and total RNA was prepared from these cells using TRIzol reagent (Invitrogen) followed by RNeasy Mini kit (Qiagen) clean-up following manufacturer’s protocol. RNA purity was confirmed using a NanoDrop Spectrophotometer (Thermo Fisher Scientific).

#### Primary cytotrophoblasts

Prior to RNA extraction, human placental chorionic villi were digested into single-cell suspension as described here for flow cytometry analysis and as in Shannon et al., 2022. Following this, single cells were stained with 7-Aminoactinomycin D (7-AAD) viability marker, EGFR BV605 (clone AY13; 1:50; Biolegend), HLA-G APC (clone MEMG-9; 1:50; Abcam), CD45 ef450 (clone 30-F11; 1:150; Invitrogen). Primary EGFR^+^, HLA-G^-^ cytotrophoblasts were isolated by flow analytical cell sorting and total RNA was prepared from these cells using TRIzol reagent (Invitrogen) followed by RNeasy Mini kit (Qiagen) clean-up following manufacturer’s protocol. RNA purity was confirmed using a NanoDrop Spectrophotometer (Thermo Fisher Scientific).

### cDNA synthesis and qPCR

100 ng RNA was reverse transcribed using qScript cDNA SuperMix synthesis kit (Quanta Biosciences) and subjected to qPCR 2^-ΔΔCT^ analysis, using PowerUP SYBR Green Master Mix (Thermo Fisher Scientific) on a StepOnePlus Real-Time PCR System (Thermo Fisher Scientific). All reactions were performed in triplicate. Forward and reverse primer sets for: *ITGA6* (F: 5’-CACTCGGGAGGACAACGTGA-3’; R: 5’-ACAGCCCTCCCGTTCTGTTG-3’), *TEAD4* (F: 5’-CGGCACCATTACCTCCAACG-3’; R: 5’-CTGCGTCATTGTCGATGGGC-3’), *ERVFRD-1* (F: 5’-ACTAGCAGCCTACCGCCATC-3’; R: 5’-TCTGGGCGAGGCTGGATAAG-3’), *CGB* (F: 5’-GCCTCATCCTTGGCGCTAGA-3’; R: 5’-TATACCTCGGGGTTGTGGGG-3’), *ITGA5* (F: 5’-GGGTCGGGGGCTTCAACTTA-3’; R: 5’-ACACTGACCCCGTCTGTTCC-3’), *NOTCH1* (F: 5’-GCCTGAATGGCGGGAAGTGT-3’; R: 5’-TAGTCTGCCACGCCTCTGC-3’), *HLA-G* (F: 5’-CCCACGCACAGACTGACAGA-3’; R: 5’-AGGTCGCAGCCAATCATCCA-3’), *ITGA1* (F: 5’-ACGTGCTGTTCATCTCGGCA-3’; R: 5’-TGCAGTTCCAAAACGAGCCC-3’), and *GAPDH* (F: 5’-AGGGCTGCTTTTAACTCTGGT-3’; R: 5’-CCCCACTTGATTTTGGAGGGA-3’). All raw data were analyzed using StepOne Software v2.3 (Thermo Fisher Scientific). Threshold cycle values were used to calculate relative gene expression levels. All values were normalized to endogenous *GAPDH* transcripts.

### Immunofluorescence microscopy

Human placental villi (GA: 11.2 weeks), PD-TOrg (P1111), and hTSC-TOrg (CT29) samples were fixed in 4% paraformaldehyde overnight at 4°C. Tissues were paraffin embedded and sectioned at 6 µm intervals onto glass slides. Immunofluorescence was performed as previously described in Shannon et. al., 2022. Briefly, *in vitro* trophoblast organoids or *in vivo* placental tissues underwent antigen retrieval by heating slides in a microwave for 5 x 2 minute intervals in citrate buffer (pH 6.0). Sections were incubated with sodium borohydride for 5 minutes at room temperature, followed by Triton X-100 permeabilization for 5 minutes at room temperature. Slides were blocked in 5% normal goat serum/0.1% saponin for 1 hour at room temperature and incubated with combinations of the indicated antibodies overnight at 4°C: mouse monoclonal anti-HLA-G (clone 4H84; 1:100; Exbio); mouse monoclonal anti-cytokeratin 7 (clone RCK105; 1:100; Santa Cruz Biotechnology), rabbit monoclonal anti-EGR1 (1:50; 15F7; Cell Signaling Technology); rabbit polyclonal anti-TEAD4 (1:200; Millipore Sigma), rabbit polyclonal anti-FOXO4 (NBP2-32614; 1:50; Novus Biologica), mouse monoclonal anti-HLA-G (clone 4H84; 1:100; Exbio), rabbit monoclonal anti-EGFR (1:50; D38B1; Cell Signaling Technology), and rabbit polyclonal IGG (1:50; eBioscience). Following overnight incubation, sections and coverslips were washed with PBS and incubated with Alexa Fluor goat anti-rabbit 568 and goat anti-mouse 488 conjugated secondary antibodies (1:200; A11036, A32723; Life Technologies) for 1 hour at room temperature, washed in PBS, and mounted using ProLong Gold mounting media containing DAPI (Life Technologies). Slides were imaged using an AxioObserver inverted microscope (Carl Zeiss) using 20x Plan-Apochromat/0.80NA or 40x Plan-Apochromat oil/1.4NA objectives (Carl Zeiss). An ApoTome .2 structured illumination device (Carl Zeiss) set at full Z-stack mode with five phase images was used for image acquisition. Patient/sample metadata are summarized in Table S2A.

### RNA Scope *in situ* hybridization

Human placental villi (6-12 weeks’ gestation; *n*=2), PD-TOrg (P1111), and hTSC-TOrg (CT29) were fixed in 2% paraformaldehyde overnight at 4°C. Tissues were paraffin embedded and sectioned at 6 µm onto glass slides. The RNAscope 2.5HD Assay - RED (ACD Bio) was performed according to manufacturer’s instructions. The Hs-Integrin alpha 2-C1-Homo sapiens mRNA (Cat#418461, ACD bio) probe was used. Slides were imaged with an AxioObserver inverted microscope (Car Zeiss, Jena, Germany) using 20X Plan-Apochromat/0.80NA or 40X Plan-Apochromat/1.4NA objectives (Carl Zeiss). Patient/sample metadata are summarized in Table S2A.

### Statistical analysis

Data are reported as median values with standard deviations. All calculations were carried out using GraphPad Prism 9 (version 9.3.1; San Diego, CA). For single comparisons, Mann-Whitney non-parametric unpaired t-tests were performed. For multiple comparisons, one-way Kruskal-Wallis ANOVA followed by Dunn’s multiple comparison test was performed. One-way ANOVA followed by Tukey post-test were performed for all other multiple comparisons. Differences were accepted as significant at *P* < 0.05. For scRNA-seq statistical analyses, please refer to the relevant scRNA-seq analysis methods sections.

## DATA AVAILABILITY/ACCESSION NUMBER

The ArrayExpress accession number for the publicly available data reported in this paper is: E-MTAB-6701. The GEO accession numbers for publicly available data reported in this paper is: GSE174481. The code used for all data processing and bioinformatic analyses are available at: https://github.com/MatthewJShannon.

## AUTHOR CONTRIBUTIONS

Conceptualization: M.J.S., A.G.B.; Methodology: M.J.S., J.W., J.B., G.L.M., B.K., B.C., M.S.P., A.G.B.; Formal analysis: bioinformatic – M.J.S., benchwork – M.J.S., J.W., G.L.M., B.K., B.C., M.S.P.; Resources: A.G.B.; Writing-original draft: M.J.S., A.G.B.; Writing – review & editing: M.J.S., J.W., J.B., G.L.M., B.K., B.C., M.S.P., W.P.R., P.C.K.L., A.G.B.; Supervision: A.G.B.; project administration: A.G.B.; Funding acquisition: P.C.K.L., A.G.B.

## ACKNOWLEDGEMENTS

The authors extend their sincere gratitude to the hard work of staff at the British Columbia Women’s Hospital CARE Program for recruiting participants into our study and the staff at the University of British Columbia Biomedical Research Centre, especially Yiwei Zhao and Tara Stach, for assistance with PD-TOrg drop-seq library sequencing. As well, we would like to thank Sana Samadi for her assistance in optimizing FOXO4 immunofluorescence.

## ABBREVIATIONS

cCTB: Column cytotrophoblast

CTB: Cytotrophoblast

DGE: Differential gene expression

EVT: Extravillous trophoblast

GA: Gestational age

HLA: Human leukocyte antigen

hTSC-TOrg: Human trophoblast stem cell line-derived trophoblast organoid

MPI: Maternal-Placental interface (*in vivo*) trophoblasts

PD-TOrg: Patient-derived trophoblast organoid

scRNA-seq: single-cell RNA sequencing

SCT: Syncytiotrophoblast

SCTp: Syncytiotrophoblast precursor

TOrg: 3D trophoblast organoid

UMAP: Uniform manifold approximation and projection

## TITLES AND LEGENDS TO FIGURES

**Figure S1:**
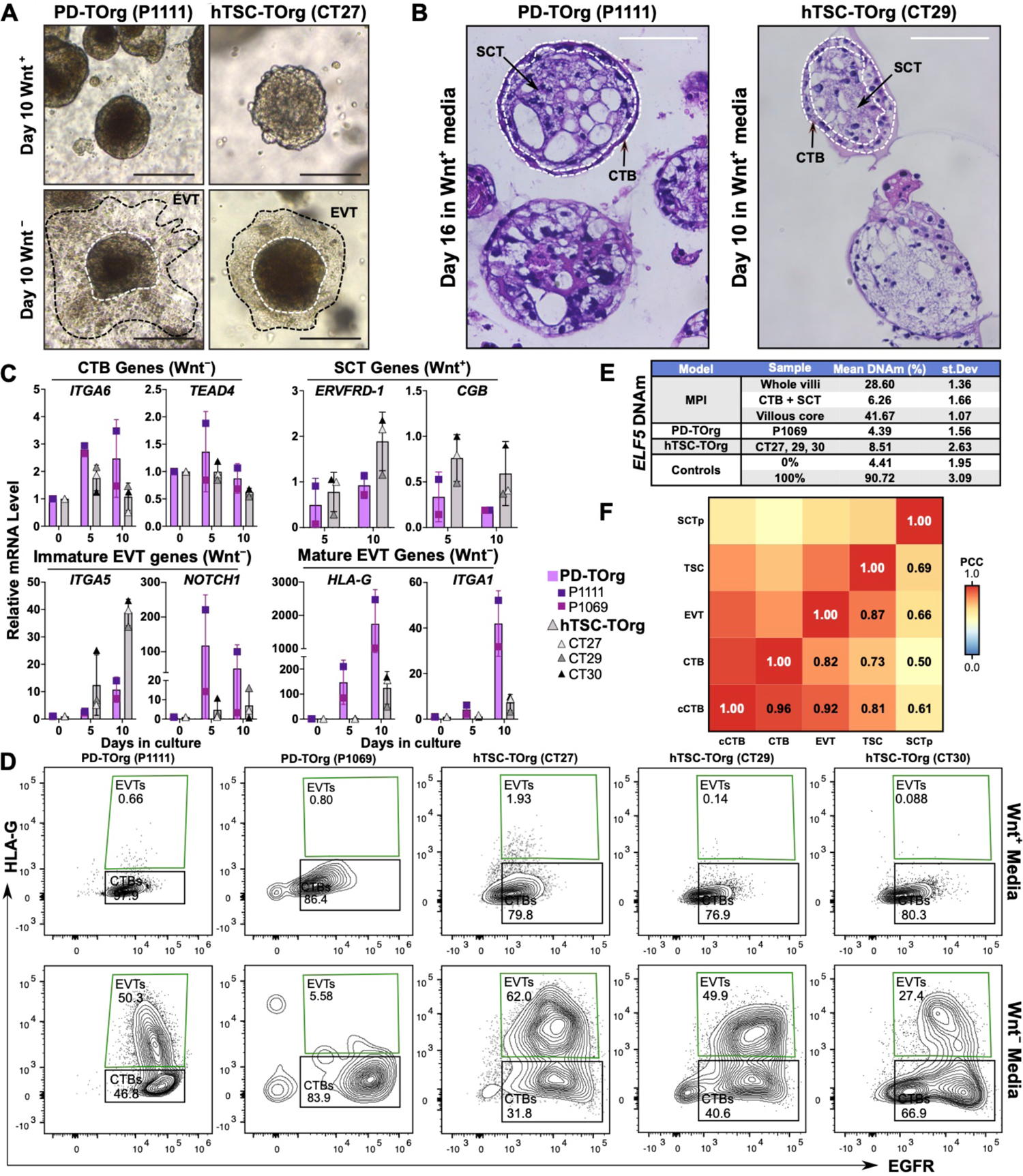
PD-TOrg model establishment. **(A)** Representative brightfield microscopy images of PD-(left) and hTSC-TOrg (right) 3D cultures following 10 days culture in regenerative (Wnt^+^) or EVT-differentiated (Wnt^-^) media. EVT outgrowth is shown by hashed line. Bar = 400 µm. **(B)** H&E staining of PD-(left) and hTSC-TOrg (right) histological sections. TOrgs were cultured to 200 μm endpoint in Wnt^+^ media prior to sectioning. Bar = 200 µm. **(C)** Relative measurement of CTB/progenitor (*ITGA6*, *TEAD4*), syncytial (*ERVFRD-1*, *CGB*), immature (*ITGA5*, *NOTCH1*), and mature EVT (*HLA-G*, *ITGA1*) mRNA levels by qPCR in PD- or hTSC-TOrgs over 10 days (D0-10) culture in regenerative (Wnt^+^) or EVT-differentiating (Wnt^-^) media. Each PD- and hTSC-TOrg sample was differentiated as one (*n*=1) replicate. **(D)** Representative flow cytometry plots showing frequencies of HLA-G^+^ and EGFR^+^ cells collected from regenerative (Wnt+) and EVT-differentiated (Wnt^-^) PD- and hTSC-TOrg cultures. Percentage of live cells are shown within EVT and CTB gates. **(E)** Table showing mean MPI, PD-, and hTSC-TOrg *ELF5* promoter methylation levels, measured by Pyrosequencing, relative to 0% and 100% methylation controls. Whole villi samples represent the entire chorionic villous while CTB + SCT samples represent the TB layers enzymatically digested from matched villous core (non-TB portions of the villous) samples using Collagenase IA. Mean data represent *n*=6 CpGs within the *ELF5* promoter/5’ UTR for each sample. Mean data represent *n*=6 CpGs within the *ELF5* promoter for each sample. **(F)** Pearson’s correlation coefficient heatmap between integrated cCTB, CTB, EVT, TSC, and SCTp trophoblast subtypes. Trophoblast subtypes were treated as pseudo-bulk samples during correlation. Abbreviations are as in Fig. 1.

**Figure S2:**
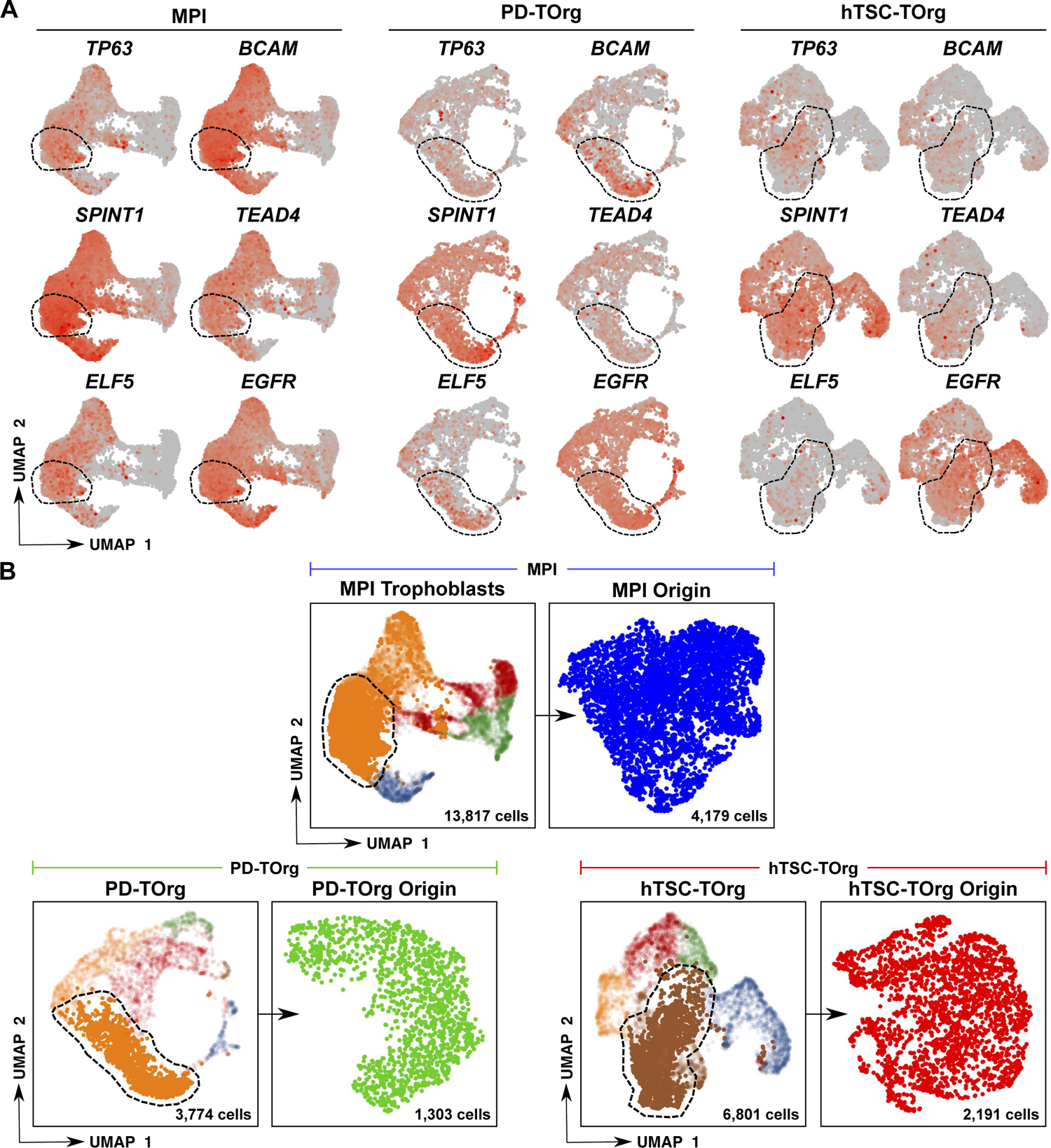
Model-specific trophoblast origin identification. **(A)** Expression (red) of progenitor CTB-associated genes plotted in MPI (left), PD- (middle), or hTSC-TOrg (right) UMAP space. Predicted origins in each model are encircled. **(B)** Model-specific trophoblast origin identification pipeline. Trophoblasts selected by the dashed line were subset for origin-specific analyses. Abbreviations are as in Fig. 1.

**Figure S3:**
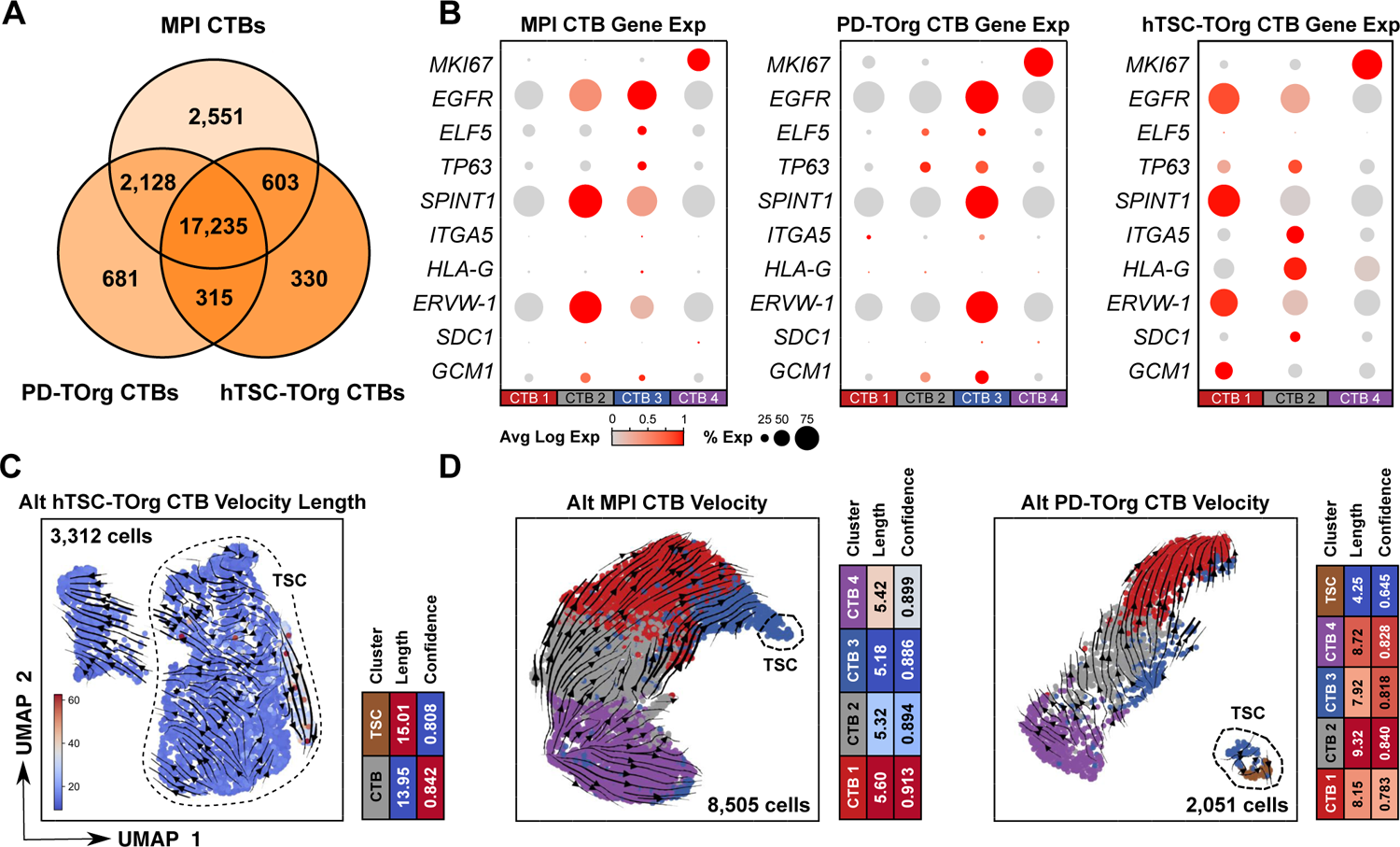
Model-specific CTB comparison. **(A)** Venn diagram comparing the bulk MPI, PD-, and hTSC-TOrg CTB transcriptomic signatures. **(B)** Dot plot showing the percent and average expression level of proliferation (*MKI67*), CTB (*EGFR*, *ELF5*, *TP63*, *SPINT1*), EVT (*ITGA5*, *HLA-G*), SCT (*ERVW-1*, *SCD1*), and TB differentiation (*GCM1*) associated genes in MPI, PD-, and hTSC-TOrg CTB. **(C)** Re-clustered hTSC-TOrg CTB and TSC UMAP. UMAP is overlain with projected RNA velocity vector field and color coordinated by velocity vector length. Associated heatmap quantifies average state vector length and confidence. **(D)** Re-clustered MPI (left) and PD-TOrg (right) CTB and TSC UMAP plot. UMAP is overlain with projected RNA velocity vector field and color coordinated by velocity vector length. Associated heatmap quantifies average state vector length and confidence. Abbreviations are as in Fig. 1.

**Figure S4:**
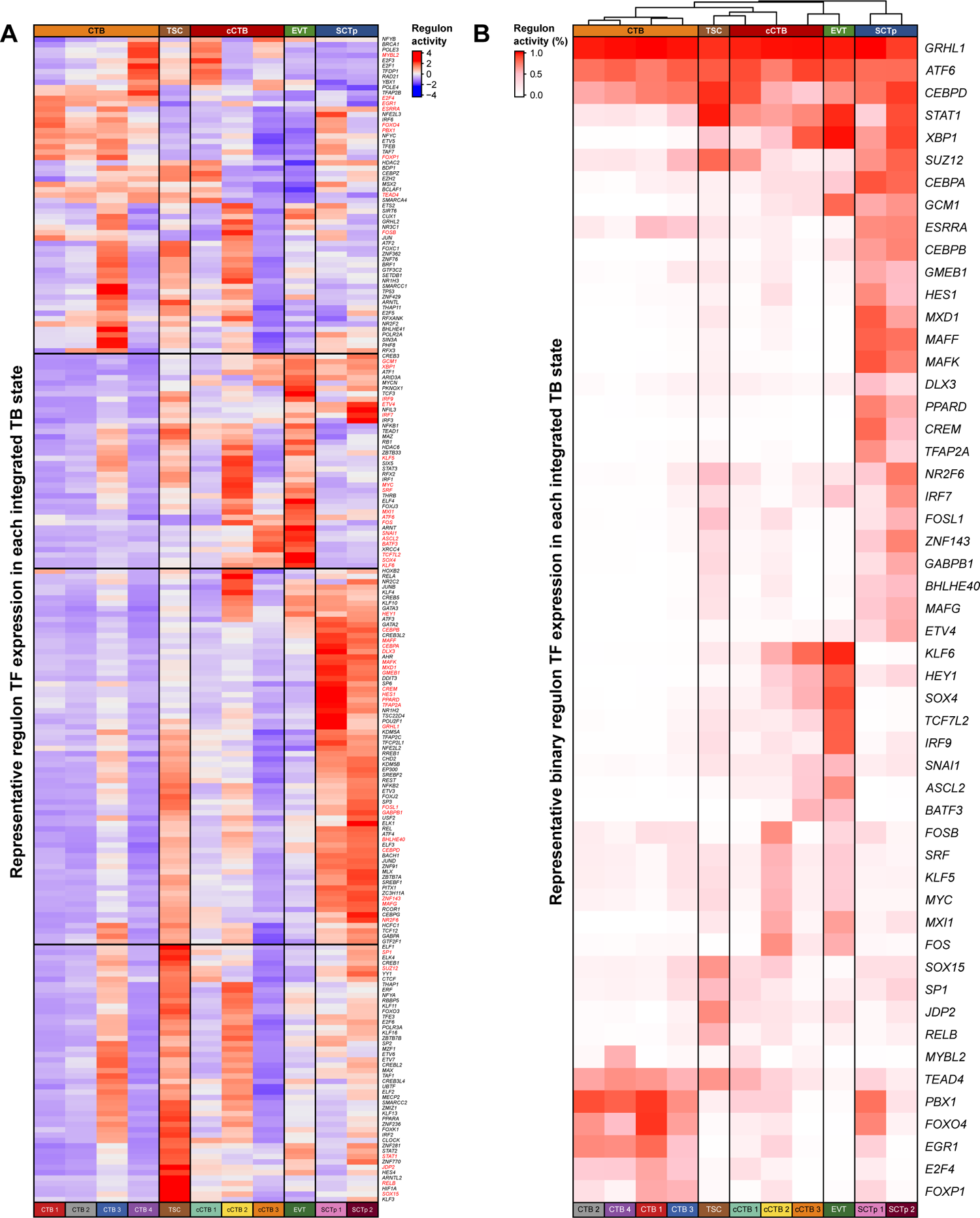
Defining trophoblast regulon specificity and regulon activity level. **(A)** Heatmap showing regulon activity across all integrated TB states from one representative SCENIC analysis. Final results were aggregated from *n*=5 independent assessments. Select genes (red) represent those plotted in panel B. **(B)** Heatmap showing percent regulon activity of significant binarized regulons across all integrated TB states from one representative SCENIC analysis. Final results were aggregated from *n*=5 independent assessments. Abbreviations are as in Fig. 1.

**Figure S5:**
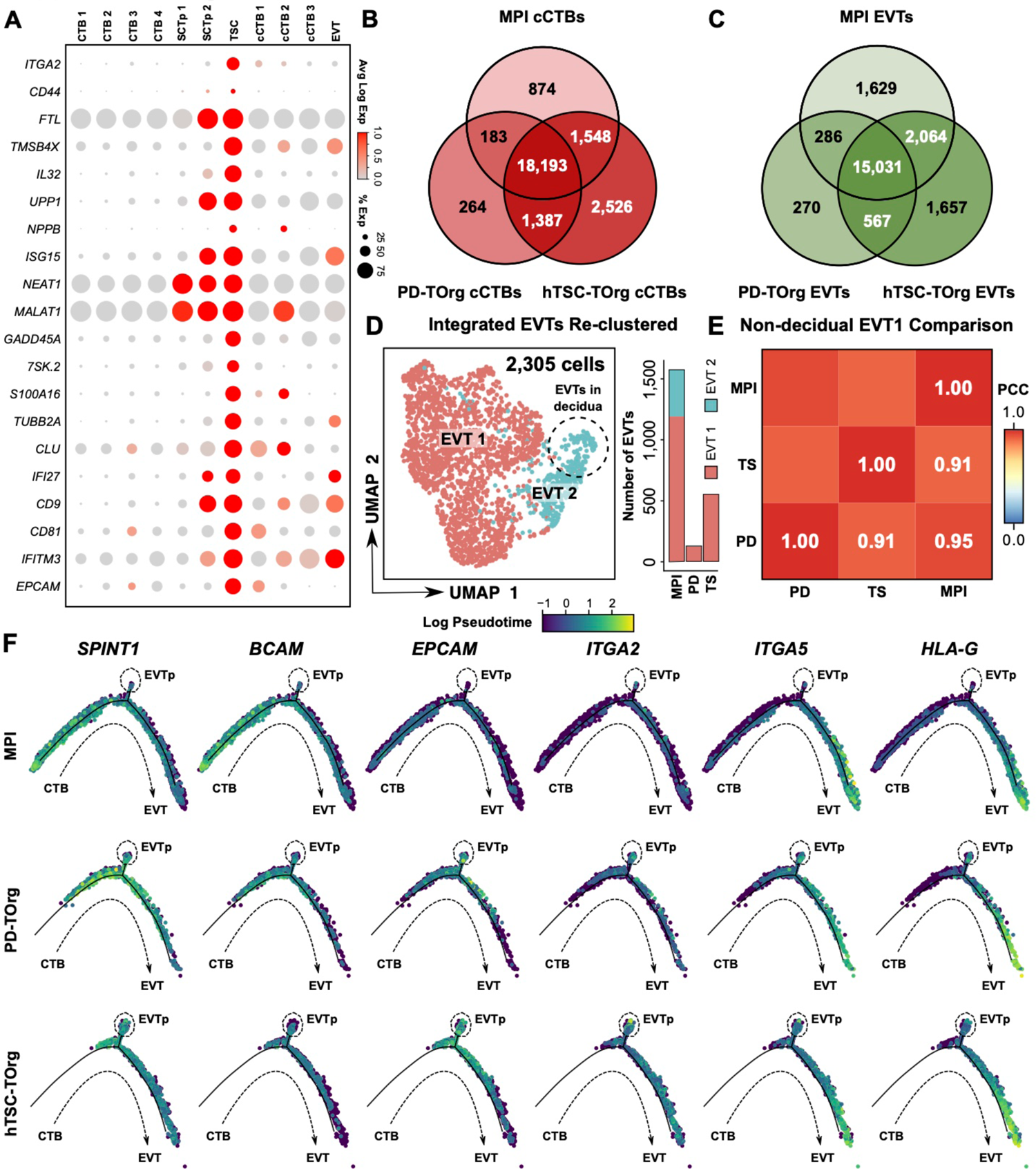
Model-specific cCTB and EVT comparison. **(A)** Dot plot showing the percent and average expression level of identified TSC enriched transcripts. **(B)** Venn diagram comparing the bulk MPI, PD-, and hTSC-TOrg cCTB transcriptomic signatures. **(C)** Venn diagram comparing the bulk MPI, PD-, and hTSC-TOrg EVT transcriptomic signatures. **(D)** Re-clustered integrated EVT UMAP plot. EVT derived from decidual tissues are encircled. Associated bar plot represents the proportion of re-clustered EVT states in each model. Abbreviations are as in Fig. 1. **(E)** Pearson’s correlation coefficient heatmap between non-decidual *in vivo* and TOrg EVT. Model-specific EVT cells were treated as pseudo-bulk samples during correlation. **(F)** Integrated Monocle2 pseudotime ordering of MPI (top), PD-(middle), and hTSC-TOrg (bottom) trophoblasts along the extravillous (towards EVT) path of differentiation (top) colour coordinated by logarithmic expression of known CTB (SPINT1, BCAM), cCTB (EPCAM, ITGA2), and EVT (ITGA5, HLA-G) gene markers across predicted pseudotime values.

**Figure S6:**
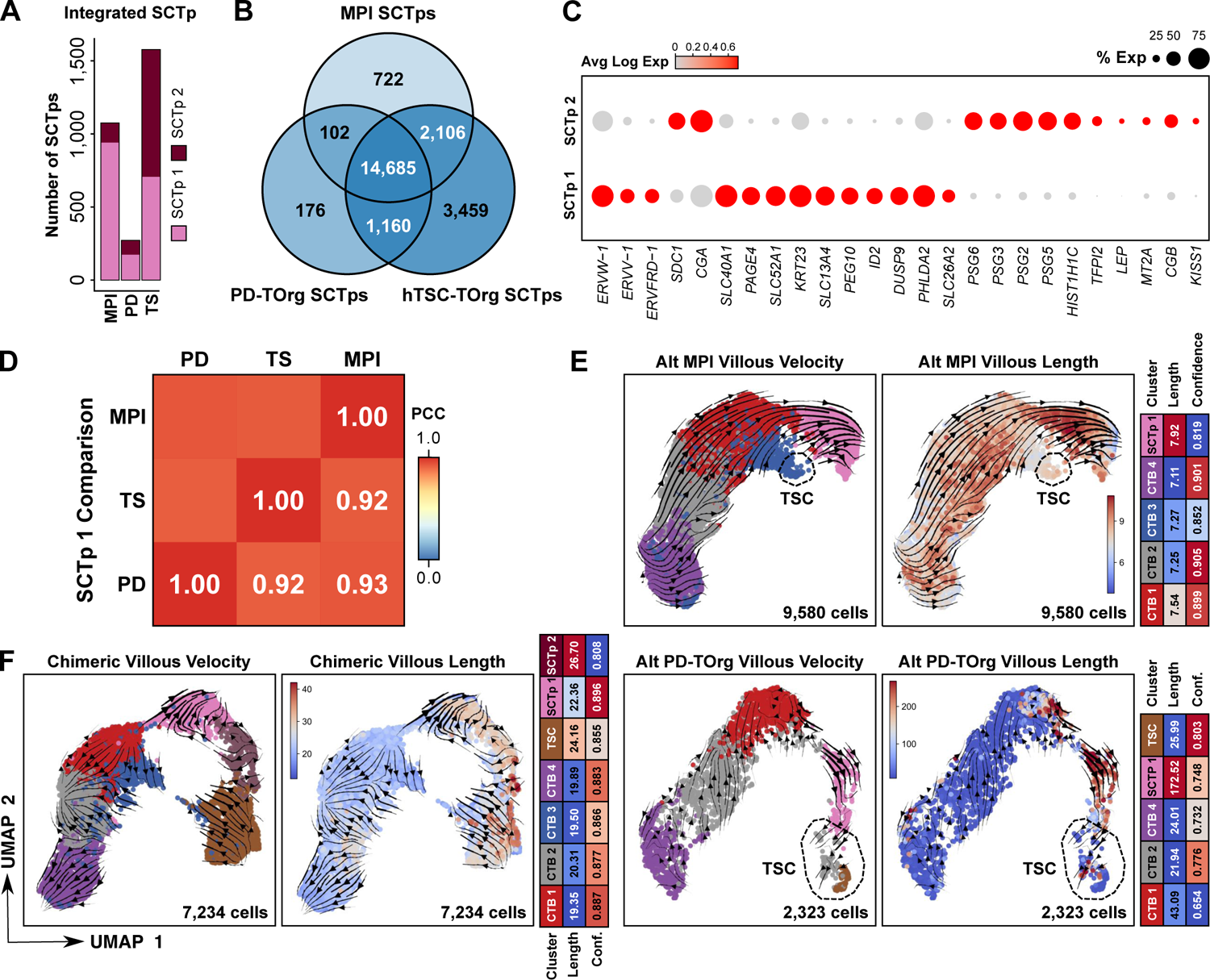
Model-specific SCTp comparison. **(A)** Bar plot showing the proportion of SCTp states in each model. **(B)** Venn diagram comparing the bulk MPI, PD-, and hTSC-TOrg SCTp transcriptomic signatures. **(C)** Dot plot showing the percent expression and average expression level of enriched integrated SCTp 1 and 2 transcripts. **(D)** Pearson’s correlation coefficient heatmap between the *in vivo* and TOrg SCTp1 state. Model-specific SCTp1 cells were treated as pseudo-bulk samples during correlation. **(E)** Re-clustered MPI (top) and PD-TOrg (bottom) CTB and TSC UMAP plots. UMAPs are overlain with projected RNA velocity vector field (left) or color coordinated by velocity vector length (right). Associated heatmaps quantify average state vector length and confidence for each model. Contributing TSCs are encircled. **(F)** Re-clustered chimeric MPI and hTSC-TOrg CTB, TSC, and SCTp UMAP plot. UMAPs are overlain with projected RNA velocity vector field (left) or color coordinated by velocity vector length (right). Associated heatmaps quantify average state vector length and confidence for each model. Abbreviations are as in Fig. 1.

