## Supplemental Figure 1 for "Single-cell assessment of trophoblast stem cell-based organoids as human placenta-modeling platforms"

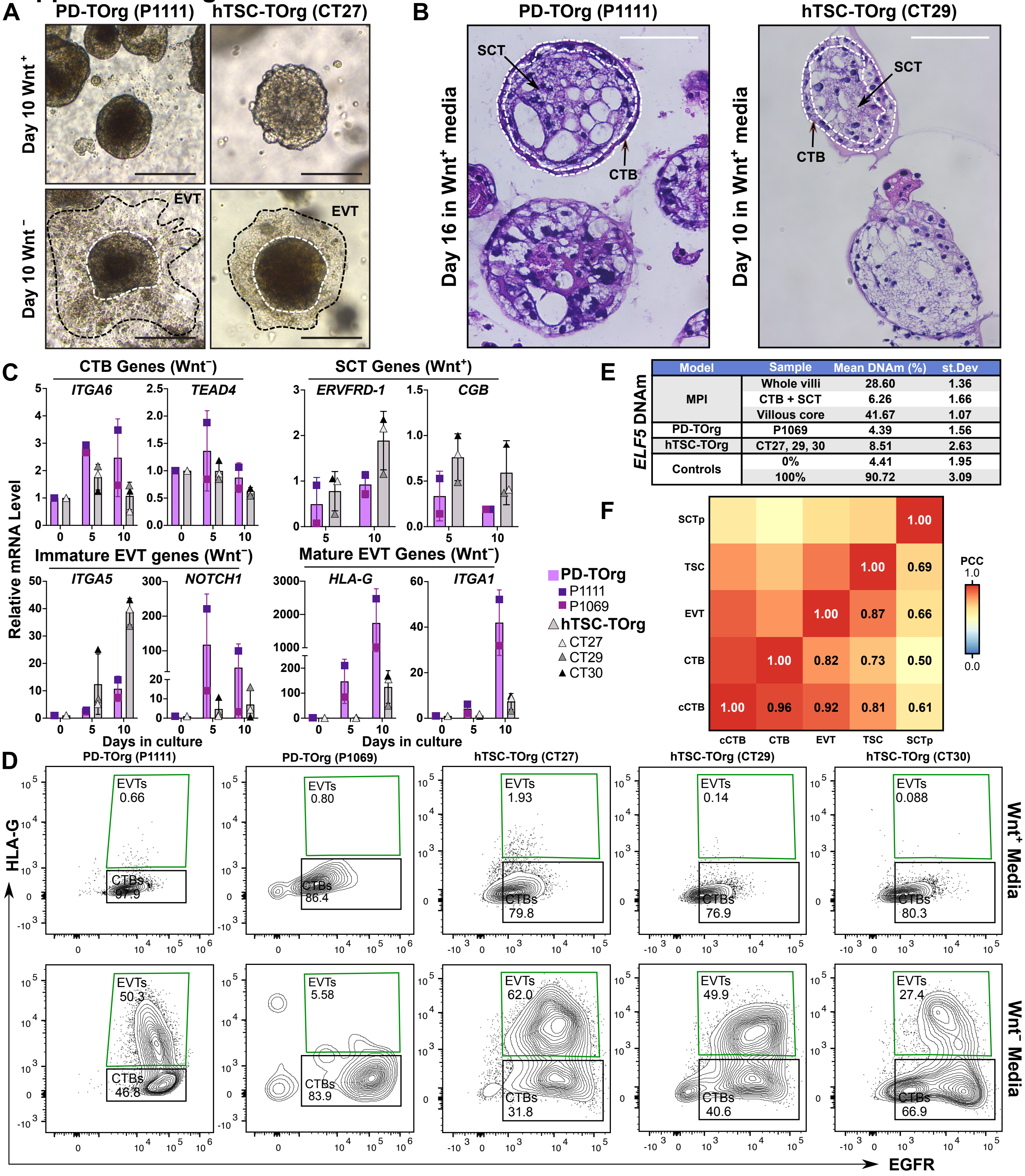


| Model | Sample | Mean DNAm (%) | st.Dev |
|---|---|---|---|
| MPI | Whole villi | 28.60 | 1.36 |
| | CTB + SCT | 6.26 | 1.66 |
| | Villous core | 41.67 | 1.07 |
| PD-TOrg | P1069 | 4.39 | 1.56 |
| hTSC-TOrg | CT27, 29, 30 | 8.51 | 2.63 |
| Controls | 0% | 4.41 | 1.95 |
| | 100% | 90.72 | 3.09 |
