## Supplementary figures and images for "Single-cell assessment of trophoblast stem cell-based organoids as human placenta-modeling platforms"

### Supplemental Figure 2

##### Supplemental Figure 2

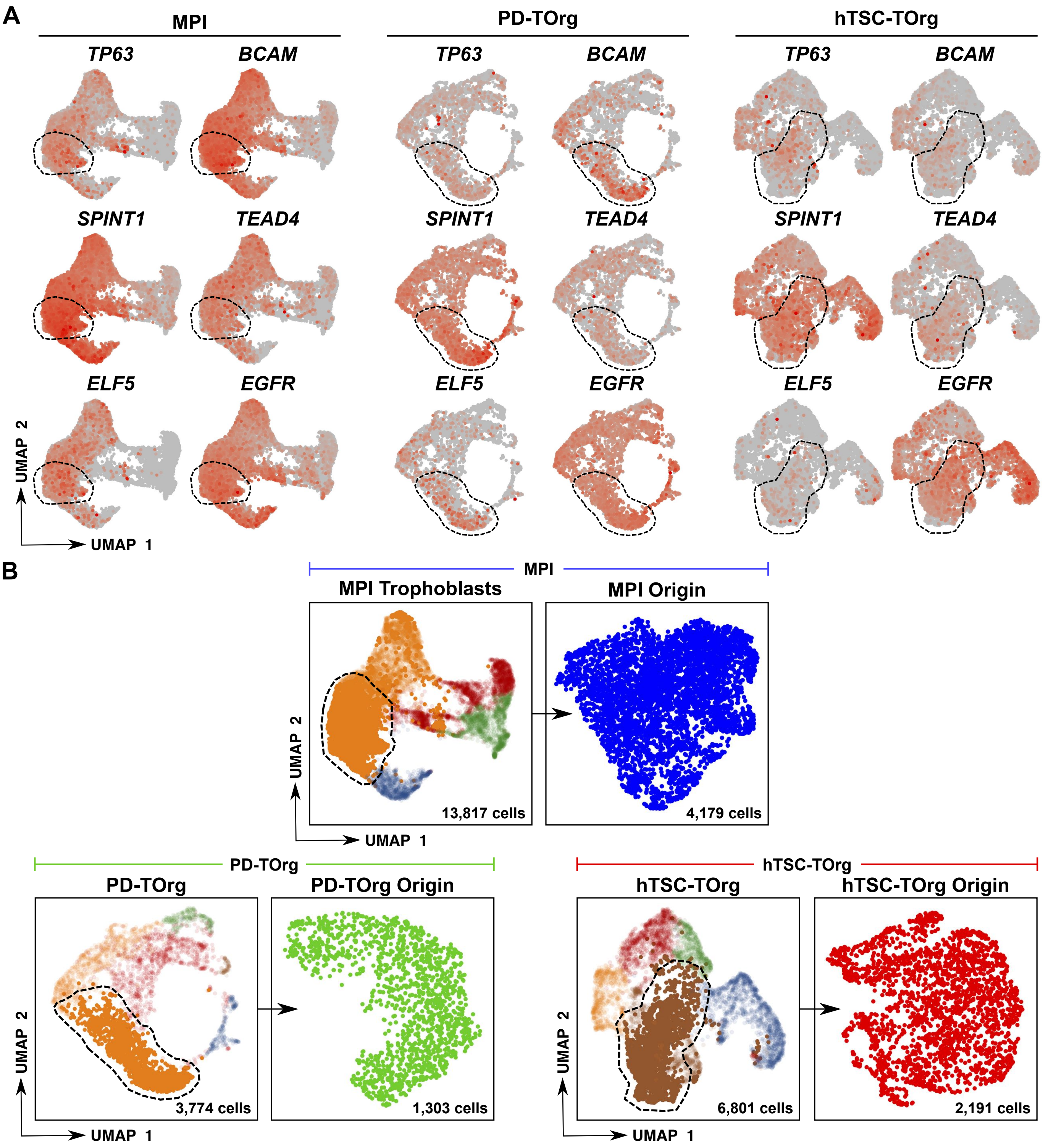

### Supplemental Figure 3

#### Supplemental Figure 3

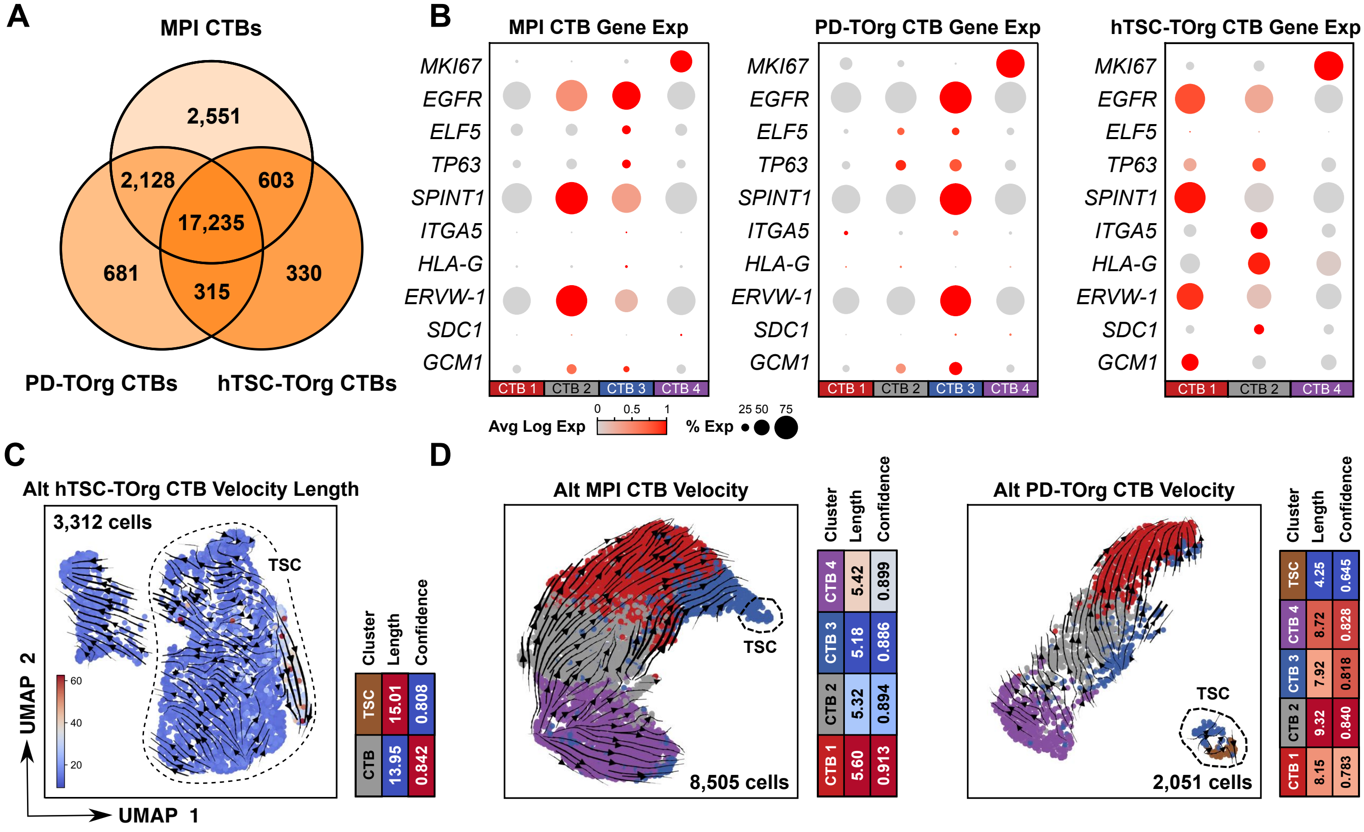

### Supplemental Figure 4

#### Supplemental Figure 4

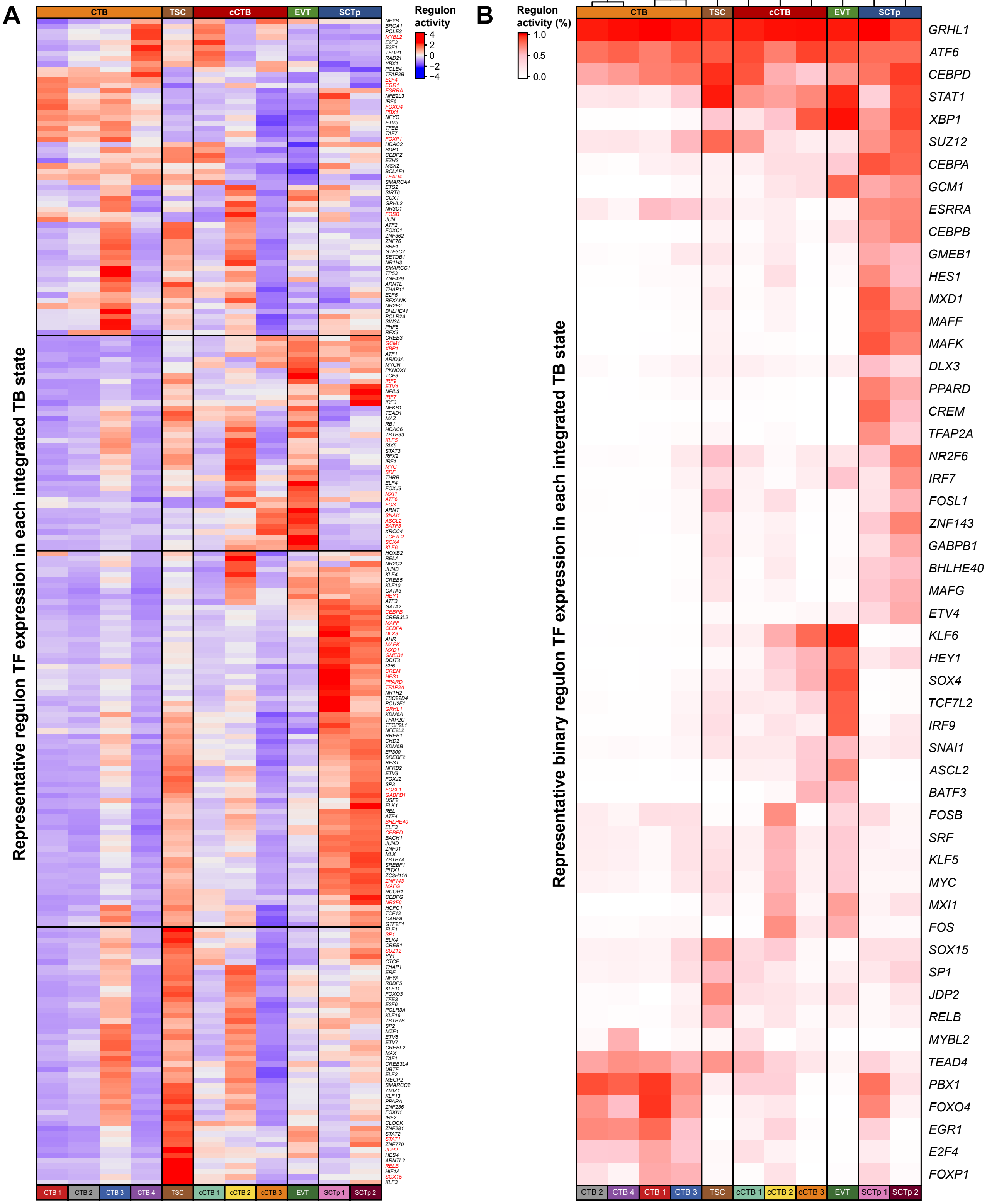

### Supplemental Figure 5

#### Supplemental Figure 5

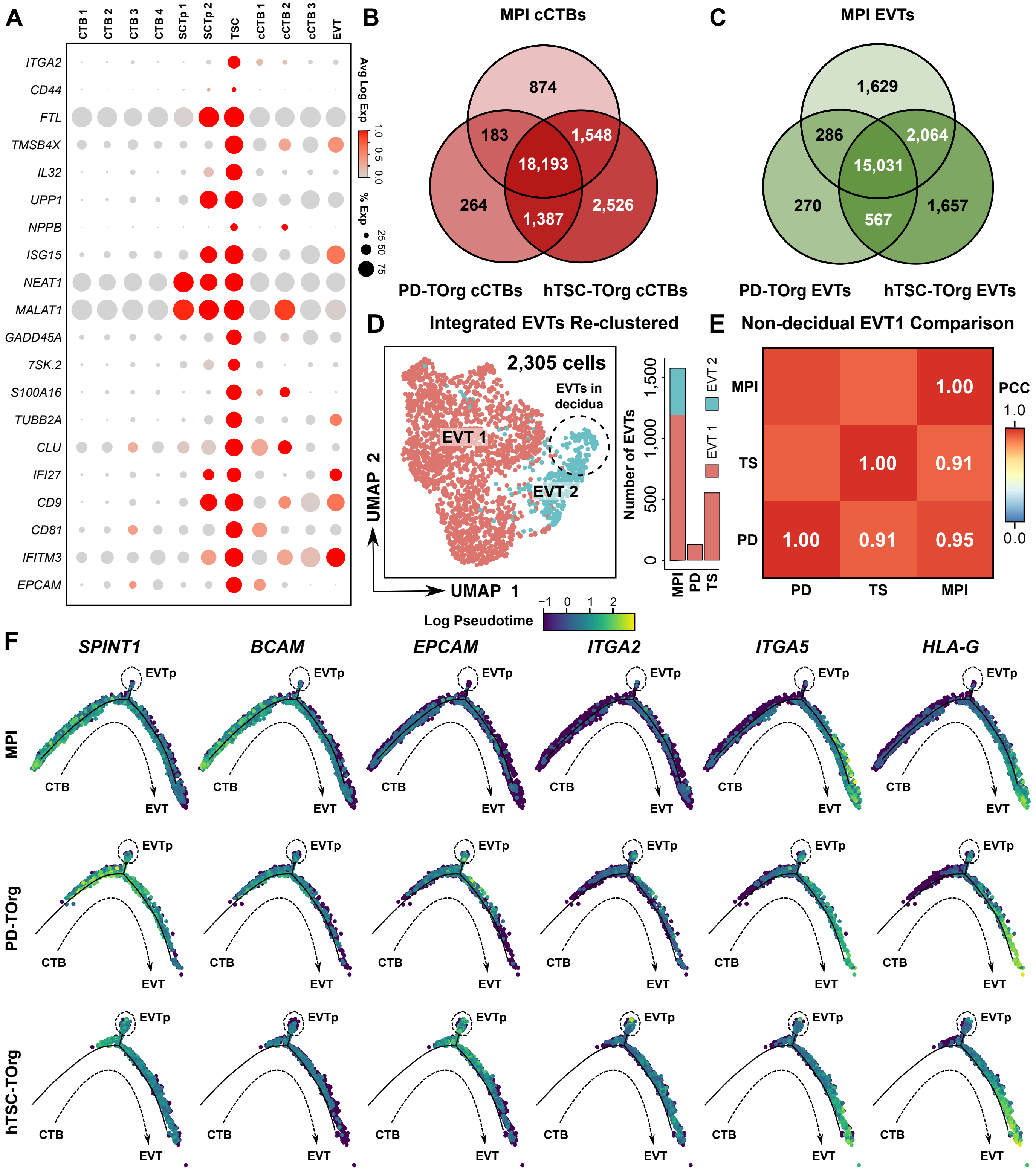

### Supplemental Figure 6

#### Supplemental Figure 6

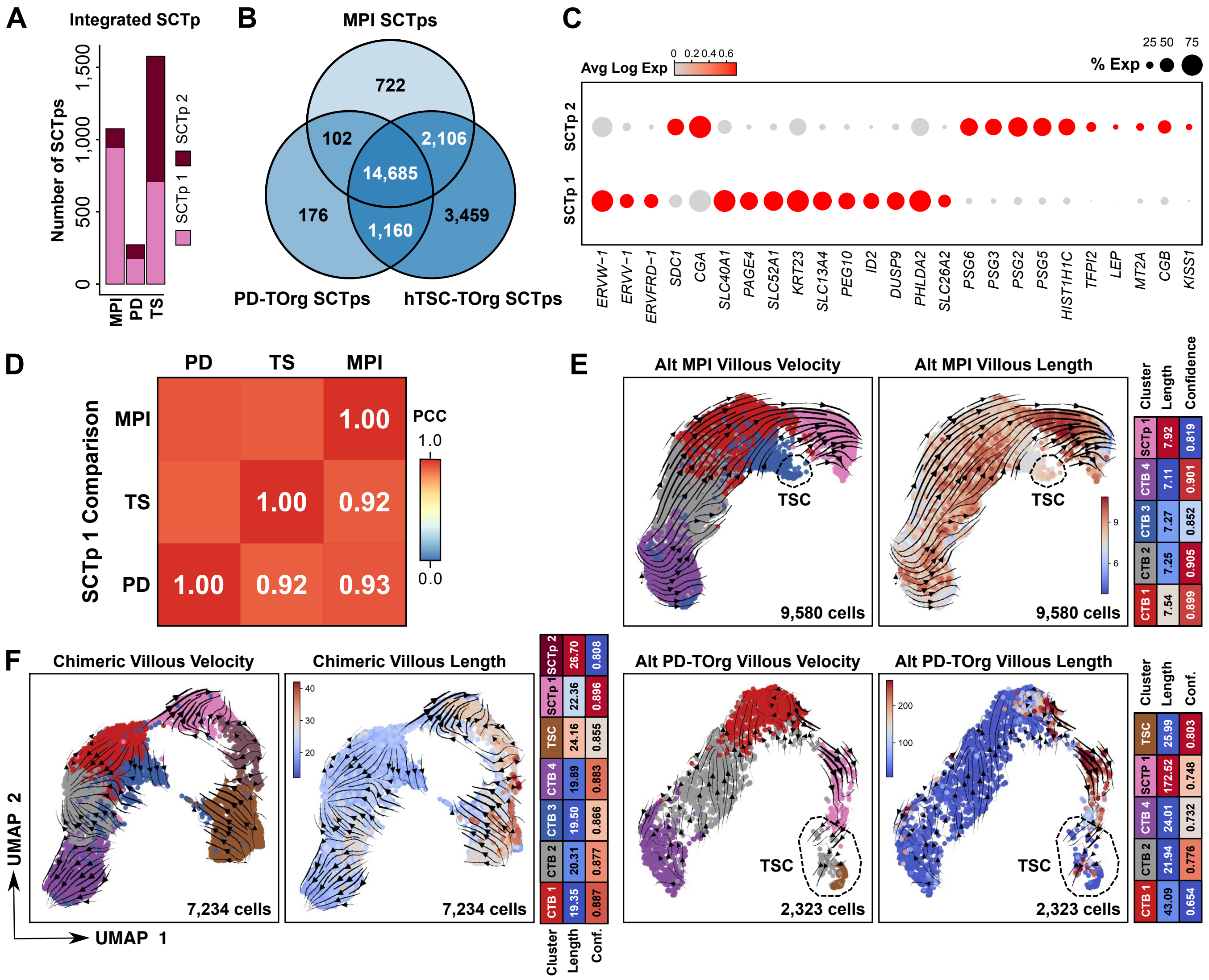
